# The initial melanoma T cell infiltrate is defined by tissue-resident programs restrained by regulatory T cells

**DOI:** 10.1101/2025.10.21.683143

**Authors:** Jason B. Williams, Shishir M. Pant, Alexander L. Kley, Bharat A. Rajmalani, Clarence Yapp, Jiang Zhang, Kyle Deans, Elizabeth Rotrosen, Angelika Sales, Peter K. Sorger, Thomas S. Kupper

## Abstract

How the immune system surveys nascent tumors and how this surveillance is subverted remain poorly understood. Using high-plex cyclic immunofluorescence and 3D imaging, we identified regulatory T (Treg) cells that co-localize with tissue-resident memory (T_RM_)-like T cells in early-stage human melanoma. In an autochthonous Braf/PTEN melanoma model expressing a defined tumor antigen, the initial CD8^+^ T cell infiltrate adopts a CD103^+^CD101^+^ T_RM_-like fate, establishing active immunosurveillance within nascent lesions. T_RM_-like cells dominate early tumors, occupy a stable epidermal niche, express effector molecules, and initiate T cell recruitment. However, Treg cells adopt a parallel tissue-resident phenotype, co-localizing with T_RM_-like cells and restraining both cytotoxic and sentinel functions. Tumor-site-specific Treg depletion reactivated T_RM_-like cells, drove robust T cell recruitment, expanded tumor-specific responses, and limited tumor growth. These findings reveal how early immunosurveillance is established through tissue-resident programs and identify Treg co-option of this response as a critical mechanism of tumor immune evasion.

**One Sentence Summary:** Nascent melanoma imprints a tissue-resident program on the initial CD8^+^ T cell infiltrate, which is suppressed by regulatory T cells as a critical checkpoint in immune evasion.

## Introduction

A major barrier to durable antitumor immunity, including responses to immune checkpoint blockade, is the emergence of resistance programs that enable tumors to escape immune pressure (*1*, *2*). While much attention has focused on T cell exhaustion and dysfunction in established tumors, the initial interactions between transformed cells and the immune system may be equally consequential. This is the period when suppressive pathways are not yet established, antigen is available, and a small number of immune cells can determine whether a nascent lesion is eliminated or progresses (*3*, *4*). Yet how nascent tumors evade elimination during this critical window remains poorly defined. Understanding how these early tumor-immune interactions unfold, and how they are subverted, is likely to reveal new therapeutic opportunities.

Tissue-resident memory T (T_RM_) cells are uniquely positioned to mediate early immunosurveillance. Unlike circulating memory T cells, T_RM_ cells remain within epithelial and mucosal tissues, poised to mount rapid, localized immune responses upon antigen encounter (*5*, *6*). They display a unique phenotype and transcriptome (*7–10*) reflecting adaptation for long-term survival and effector function in their tissue of residence. This physiology enables T_RM_ cells to act as sentinels, responding swiftly to pathogenic insults and playing a critical role in protective immunity (*11*, *12*). Mechanistically, T_RM_ cells contribute to tumor immunity through direct cytotoxicity, secretion of inflammatory cytokines, and recruitment of additional immune effector cells (*13*, *14*), functions that could be decisive during the earliest stages of tumor development.

Tissue retention by T_RM_ cells is maintained by the expression of adhesion molecules and by reduced responsiveness to cytokines and chemokines that drive recirculation. In the skin, the αE integrin CD103 (Itgae), which pairs with β7 to bind E-cadherin, and CD69, which binds and internalizes the S1P receptor (*15*), are critical for tissue retention and T_RM_ cell development (*16*). Most CD8^+^ T_RM_ cells in the skin express CD69 and/or CD103, particularly in the epidermis, and T cells bearing these markers are transcriptionally distinct from other memory T cell subsets (*7*, *8*, *16*, *17*). Cells expressing T_RM_-associated markers or a T_RM_-like transcriptional program have been identified in many cancer types (*18–22*). However, the use of CD103 and CD69 to define T_RM_ cells is imperfect: T cells upregulate CD69 following TCR stimulation and TGF-β can transiently induce CD103 expression (*10*). As a result, in chronic inflammation or progressing tumors, it can be challenging to determine whether T cells expressing T_RM_-associated markers are truly resident memory. For this reason, such cells within tumors are often referred to as “T_RM_-like” cells.

T_RM_-like cells have emerged as key players in cancer immunosurveillance and immunotherapy response. Their presence within tumors has been associated with improved prognosis and favorable responses to checkpoint blockade in multiple cancer types, including melanoma (*18*, *23–27*). Preclinical studies have demonstrated that prophylactic induction of T_RM_ cells, either via vaccination or rejection of a primary tumor challenge, confers potent protection against subsequent tumor challenges (*13*, *28*). In addition, T_RM_ cells can control primary tumor growth in a state of cancer-immune equilibrium (*29*). However, these studies have focused either on T_RM_-like populations in established tumors, where exhaustion and chronic antigen exposure obscure early differentiation events, or on T_RM_ cells seeded prophylactically before tumor challenge, which models pre-existing memory rather than natural tumor development. How T_RM_-like programs emerge and are regulated during ongoing, naturally arising tumor growth, and whether these early responses are suppressed before they can mediate rejection, remains largely unknown.

In this study, we combine spatial analysis of human melanoma with mechanistic interrogation in an autochthonous mouse model to define the fate and regulation of the earliest tumor-infiltrating CD8^+^ T cells. Using high-plex cyclic immunofluorescence and three-dimensional imaging of stage II human melanoma, we identify CD8^+^ T cells with T_RM_-like features in precursor and early invasive regions, where they localize near melanoma cells and are frequently juxtaposed with regulatory T (Treg) cells and dendritic cells. These observations suggest that tissue-adapted CD8^+^ T cells participate in tumor immunosurveillance from the earliest stages of melanoma development and may be subject to local immunoregulatory control.

To mechanistically dissect these early interactions, we employed an inducible Braf/PTEN melanoma model engineered to express ovalbumin (OVA) as a defined tumor antigen. In this autochthonous setting, the initial CD8^+^ T cell infiltrate rapidly acquires T_RM_-like characteristics, occupies a stable niche in the upper skin, and constitutes the majority of CD8^+^ tumor-infiltrating lymphocytes during early tumor development. Despite their early dominance and effector potential, these cells are rapidly restrained by co-infiltrating Treg cells, which suppress both cytotoxic activity and immune-recruitment (“sentinel”) functions. Tumor-site-specific Treg depletion reactivates these early CD8^+^ T cells, drives robust secondary T cell recruitment, and limits tumor growth. Together, these findings identify Treg-mediated suppression of the earliest CD8^+^ T cell infiltrate as a critical early step in melanoma immune evasion.

## Results

### Stage II human melanoma contains T_RM_-like cells

Given their prognostic significance, we investigated the abundance and intra-tissue distribution of T_RM_ cells in primary human melanomas. We used multiplex tissue-based imaging (cyclic immunofluorescence; t-CyCIF) and a hierarchical gating strategy (**Figure S1A-S1D**) to distinguish immune cells with related lineages in 12 stage II primary human melanoma specimens (**Figure 1A**). CD8 and CD103 co-expression was used to identify T_RM_-like CD8^+^ TILs. However, CD103 was also expressed by Foxp3^+^ regulatory T (Treg) cells, Foxp3^−^ conventional T (Tconv) cells, and dendritic cells (DCs) (**Figure S1B**). Analysis of immune cell composition revealed that most tumor samples contained T_RM_-like cells (**Figure 1B** and **S1E**), but their abundance varied substantially with the tumor, ranging from near absence (in MEL76 and MEL78) to the majority of the CD8^+^ TIL compartment (MEL71 and MEL72). In addition, CD4^+^ Foxp3^+^ T (Treg) regulatory cells were found in all tumors and composed a dominant fraction in some tumors (MEL81, MEL84, and MEL85) but not all (**Figure 1B**).

**Fig. 1.**
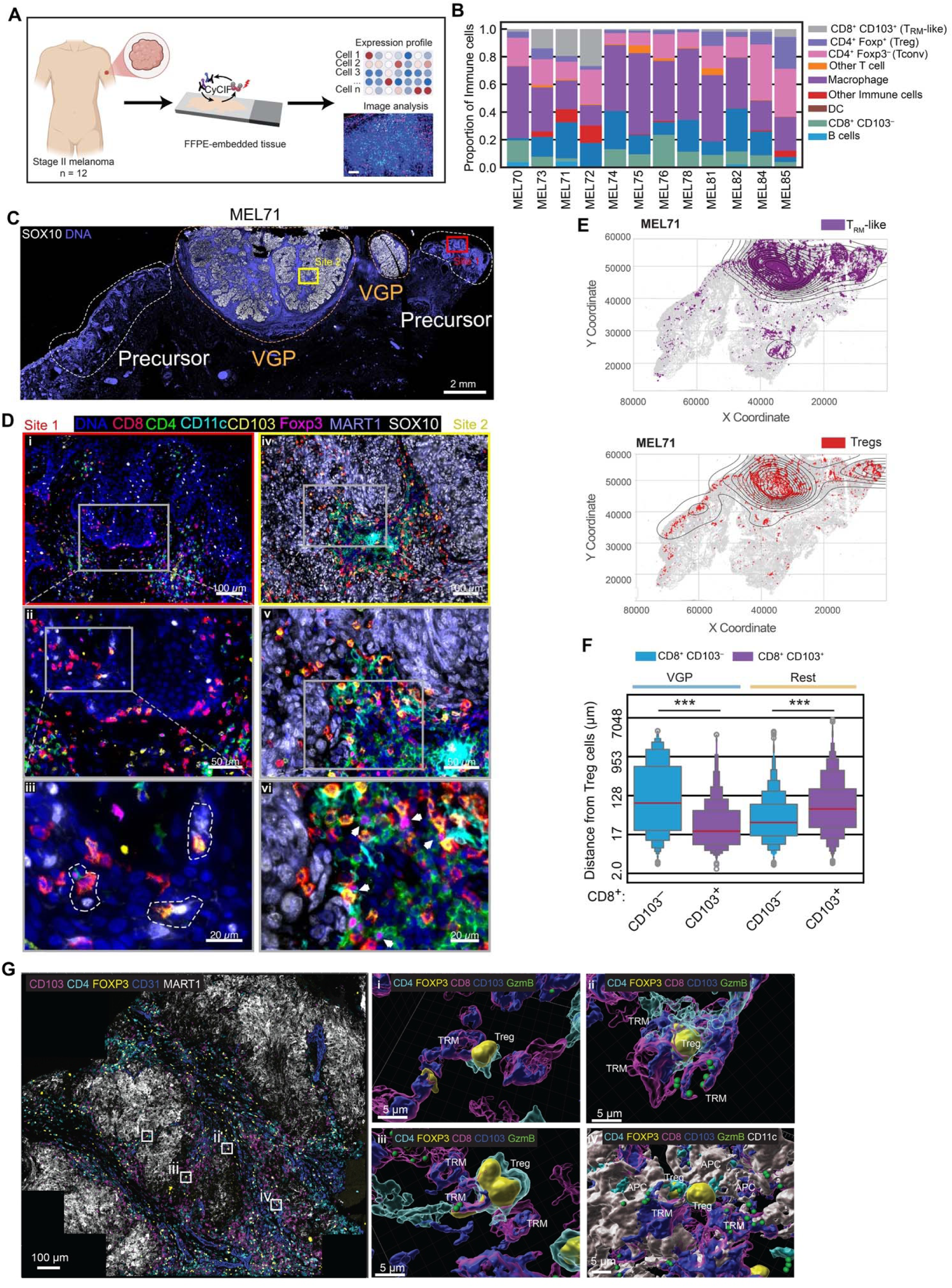
Spatial organization of T_RM_-like CD8^+^ T cells and regulatory T cells in human stage II melanoma. **A**, Schematic of the study design. FFPE-embedded stage II melanoma samples (n=12) underwent tissue-based Cyclic Immunofluorescence to generate cell phenotype and spatial data. **B**, Immune cell composition across 12 stage II melanoma samples. **C**, CyCIF image of an example specimen (MEL71) annotated with clinical histological stages: Precursor (white dotted line) and vertical growth phase (VGP, orange dotted line) showing SOX10 staining (white). Two sites were selected for zoom in view are marked with red and yellow squares. Scale bars 2 mm. **D**, Higher magnification view of the boxed regions in **C**. Site 1 (insets i-iii) highlight CD8^+^ T_RM_-like cells in the red highlighted precursor region. Dotted line in iii highlights T_RM_-like cells in proximity to MART1^+^ SOX10^+^ melanoma cells. Site 2 highlights invading CD8^+^ T_RM_-like cells in proximity to Treg cells (white arrows). **E**, Dots plots with density contours showing Treg and CD8^+^ T_RM_-like cell distribution in MEL71. **F**, Distance from Tregs to the closest CD8^+^ CD103^+^ T_RM_-like or CD103^−^ cells in VGP versus non-VGP areas across all tumor samples. **G**, 3D CyCIF image of melanoma tumor in vertical growth phase showing tumor cells (MART1: White), endothelial cells (CD31: Blue), and Immune cells (CD4: Cyan; FOXP3: Yellow; CD103: Magenta). Selected ROIs for higher magnification view are marked with white squares. Scale bars = 100 µm. Rendering of 3D CyCIF image (insets i-iv) from selected ROIs showing interaction between CD8^+^ T_RM_-like cells, Treg cells, and DCs. Scale bars = 5 µm. Significance was determined by Mann-Whitney U rank test in **F**.

Compared to their CD103^−^ counterparts, a greater fraction of T_RM_-like CD8^+^ T cells expressed the co-inhibitory receptors PD-1, LAG-3, and Tim-3 (**Figure S1B**). T_RM_-like cells were distributed in a diffuse manner near fields of melanocytic atypia (precursor regions) or in the epidermis (**Figure 1D**, site 1) but were concentrated in invasive vertical growth phase (VGP) regions (**Figure 1D**, site 2, and **Figure 1E**). In precursor regions, T_RM_-like cells could be found in close proximity to MART1^+^ SOX10^+^ melanoma cells (**Figure 1D**, inset **iii** circled regions) and were found infiltrating the tumor bed in VGP areas (**Figure 1D**, insets **iv-vi**). Treg cells exhibited a similar distribution and could be found in proximity or interacting with T_RM_-like cells in VGP areas (**Figure 1D**, inset **vi** white arrows). Cell proximity measurements revealed that within VGP regions, T_RM_-like cells were in closer proximity to Treg cells compared to their CD103^−^counterparts (**Figure 1F**), suggesting potential crosstalk between these two populations. Three-dimensional tissue images (3D CyCIF) of thick sections provided a more detailed view of cell-cell interactions and, unlike imaging of standard 5 μm sections, include many intact cells. A 35 μm thick 3D CyCIF reconstruction of invasive primary melanoma (*30*) revealed multiple instances of direct T_RM_-like:Treg cell interactions (**Figure 1G**). DCs were also frequently observed to form a triad with Treg and T_RM_-like cells (**Figure 1G** inset **iv** and **Figure 1D** inset **vi,** circled regions). Given these observations in human tumors, we sought to develop a preclinical model that could generate T_RM_-like CD8^+^ T cell populations in a melanoma TME and enable mechanistic analysis of their regulation and function including an exploration of Treg-mediated suppression of T_RM_-like cells.

### The autochthonous Braf/PTEN tumor model contains T_RM_-like CD8^+^ TIL

We evaluated several mouse tumor models to determine their capacity to generate T_RM_-like CD8^+^ TILs. In the engraftable tumor models (B16.OVA, MC38.OVA, Yumm1.7), only a small percentage of CD8^+^ TILs expressed the T_RM_ marker CD103 following subcutaneous or intradermal injection. Even when Yumm1.7 was implanted epicutaneously (*29*), the results were similar (**Figure 2A**). We then tested the genetically engineered BRaf^CA/+^/PTEN^loxp^/Tyr;;CreER^T2^ (BP) mouse model, which better replicates the microenvironmental cues due to its autochthonous origin. BP tumors harbored a distinct population of CD103^+^ CD8^+^ TILs 40–50 days post 4-hydroxytamoxifen (4-OHT) induction (**Figure 2A**). To facilitate tracking of antitumor immune responses, we introduced an EGFPOVA fusion protein into the BP model by crossing to Rosa26-Stop-flox-EGFPOVA (*31*) mice, generating BRaf^CA/+^/PTEN^loxp^/R26^EGFPOVA^/TyrCreER^T2^ (BPO) mice (**Figure 2B**). BPO tumors begin as hyperpigmented foci that gradually grow in size and ultimately merge into a single larger tumor (**Figure 2C**). Flow cytometric analysis confirmed enrichment of CD103^+^ CD8^+^ TILs within BPO tumors (**Figure 2D**), with higher CD69 expression in CD103^+^ CD8^+^ TILs compared to their CD103^−^ counterparts (**Figure 2E**). To assess whether these T_RM_-like cells arose as a function of the autochthonous environment, we established BPO-derived tumor cell lines and engrafted them into syngeneic mice. Engrafted tumors exhibited significantly lower frequencies of CD103^+^ CD8^+^ TILs compared to primary autochthonous tumors (**Figure S2A** and **S2B**), suggesting that factors intrinsic to the native skin microenvironment contribute to CD8^+^ T_RM_-like T cell development.

**Fig. 2.**
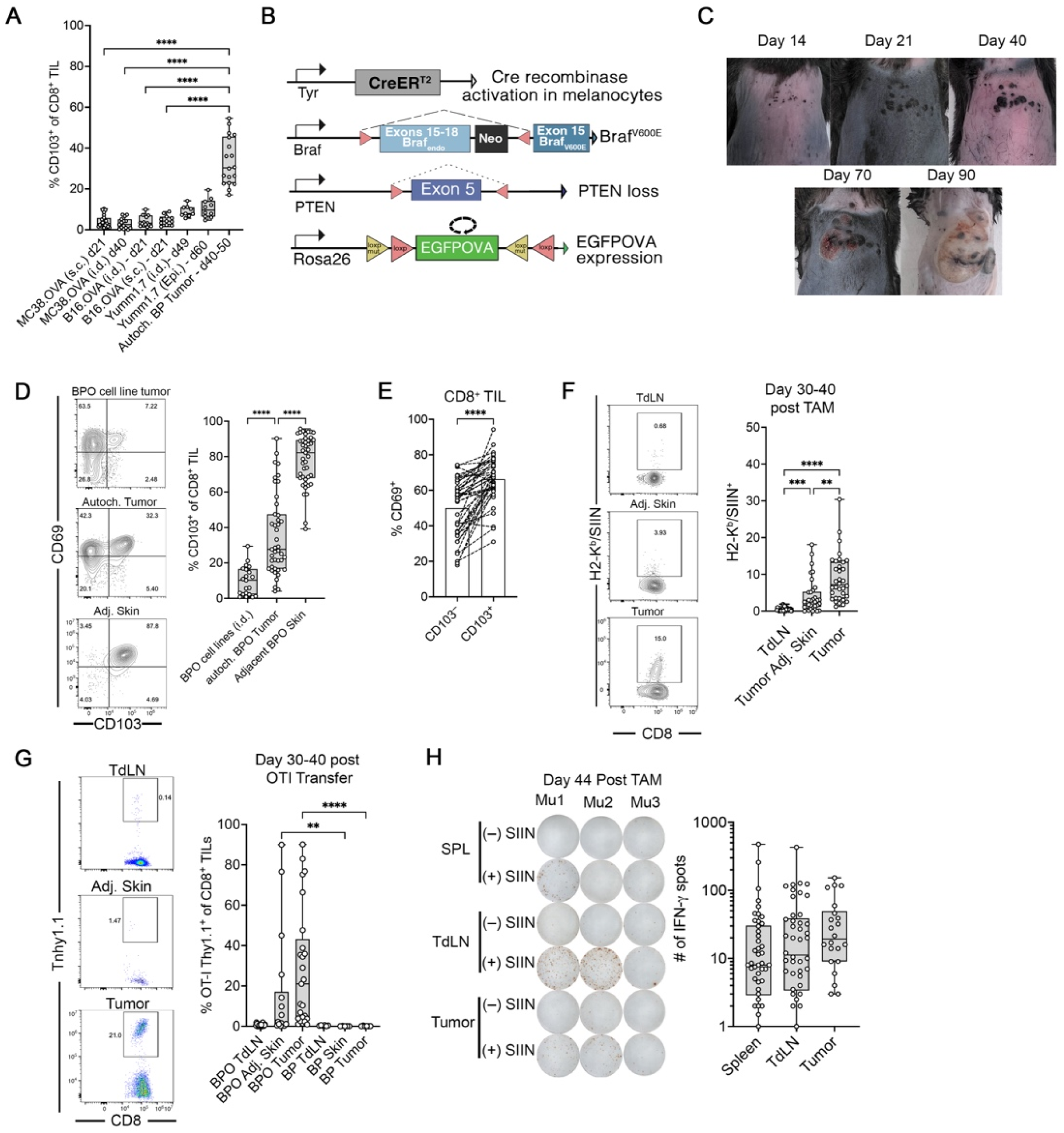
T_RM_-like CD8^+^ TILs arise in the autochthonous BPO melanoma model. **A**, Frequency of CD103^+^ CD8^+^ TILs in engraftable (B16, MC38, Yumm1.7) versus autochthonous BP tumors. **B,** Schematic of BPO model generation (BP x Rosa26-EGFPOVA). **C,** Images of BPO tumor progression. **D,** Representative flow cytometry plots for CD103 and CD69 on CD8^+^ TILs from BPO tumors and tumors from i.d. engrafted BPO-derived cell lines. **E**, comparison of CD69 expression on CD103^+^ versus CD103^−^ CD8^+^ TILs. **F**, H2-K^b^/SIINFEKL pentamer^+^ CD8^+^ T cell frequencies in TdLN, tumor-adjacent skin and tumor. **G**, OT-I transfer frequencies in TdLN, tumor adjacent skin, and tumor in BPO and BP mice. **H**, IFN-γ ELISPOT against SIINFEKL in the spleen, TdLN, and tumor cells from BPO mice. Data in panels **A** and **D-H**, display all mice from 2-5 independent experiments. Significance was determined by a Kruskal-Wallis test in **A, D, F-H**, and a Wilcoxon matched-pair signed rank test in **E**. ^+^

We next tested whether tumor-specific immune responses could be detected in the BPO model. Using H2-K^b^/SIINFEKL-pentamer staining, we identified endogenous OVA-specific CD8^+^ T cells in the tumor, tumor-draining lymph node (TdLN), and adjacent skin 30–40 days after tumor induction – with highest frequencies in the tumor (**Figure 2F**). Adoptive transfer of OVA-specific, CD8^+^ T cells (OT-I T cells) also resulted in recruitment of OT-I T cells to the tumor, which was not observed in BP (EGFPOVA-negative) control tumors (**Figure 2G**). OT-I cells also proliferated robustly in TdLNs of tumor-bearing mice, but not in tumor-free Braf/PTEN/EGFPOVA mice, demonstrating tumor-specific priming (**Figure S2C**). Finally, IFN-γ ELISPOT assays confirmed systemic tumor-specific CD8^+^ T cell responses in the spleen, TdLN, and tumor (**Figure 2H**). Collectively, these findings establish the BPO model as a robust system to study tumor-specific T_RM_-like CD8^+^ T cell responses within an autochthonous melanoma setting.

### T_RM_-like CD8^+^ TILs are phenotypically and transcriptionally distinct

With the BPO model established, we next characterized the T_RM_-like CD8^+^ TIL by bulk RNASeq of CD103^+^ CD69^+^ CD8^+^ (DP: double positive) and CD103^−^ CD69^−^ CD8^+^ (DN: double negative) TILs. We compared DP and DN populations from the tumor to DP cells from tumor adjacent skin. As an additional T_RM_ cell control, we isolated DP cells from the skin of mice that had rejected epicutaneous Yumm1.7_OVA_ tumors (*29*). Differentially expressed genes were determined by normalizing to CD8 T cells from the spleen of naïve mice. We found 722 differentially expressed genes. A shared core of genes across all groups (gene set 1) included transcription factors (*Bhlhe40*, *Irf5*, *Rora*, *Zeb2,* and *Ikzf2),* cell surface receptors (*Havcr2*, *Pdcd1*, *Cd244a*, *Klrk1, Fcer1g, Entpd1*), chemokines (*Ccl3 and Ccl4*), and effector molecules (*Ifng* and *Gzmb*), all of which are consistent with activated T cell effector functions. Genes involved in lipid metabolism, including Fabp4, CD36, and Cav1, were selectively upregulated in skin-derived T_RM_ cells (gene set 2), reflecting metabolic adaptations to epithelial niches, and transcriptional regulators (*Zfp36*, *Crem*, *Runx2*, *Socs3*, *Dusp5*, *Klf6*, and *Fosl2*) were upregulated solely in the tumor-rejected skin T_RM_ group. In contrast, tumor-associated samples (gene sets 3 and 4) contained upregulated inflammation-related genes (Tnfrsf9, Fasl, Ccr5) and purinergic pathway genes (CD38, P2rx7).

Next, we searched for genes that distinguish T_RM_-like cells from other T cells by comparing the DP and DN populations (group 5). As expected, *Itgae* was upregulated in the DP groups, along with other T_RM_-associated genes (*32*) (*Jun*, *Ccr10*, *Gzmc*, *Hspa1b*, *Hspa1a*, *Atf3*) and the cell surface receptor *Cd101*. Genes upregulated in all groups except for DN cells in tumor adjacent skin (group6) included *Cxcr6, Ccr8, Xcl1, Tnf, Nr4a2*, and *Tox2.* Finally, downregulated genes in all groups included those involved in naïve and circulating memory T cell stemness and maintenance included *Klf2, Ccr7, Sell,* and *S1pr1* (**Figure 3A**).

**Fig. 3.**
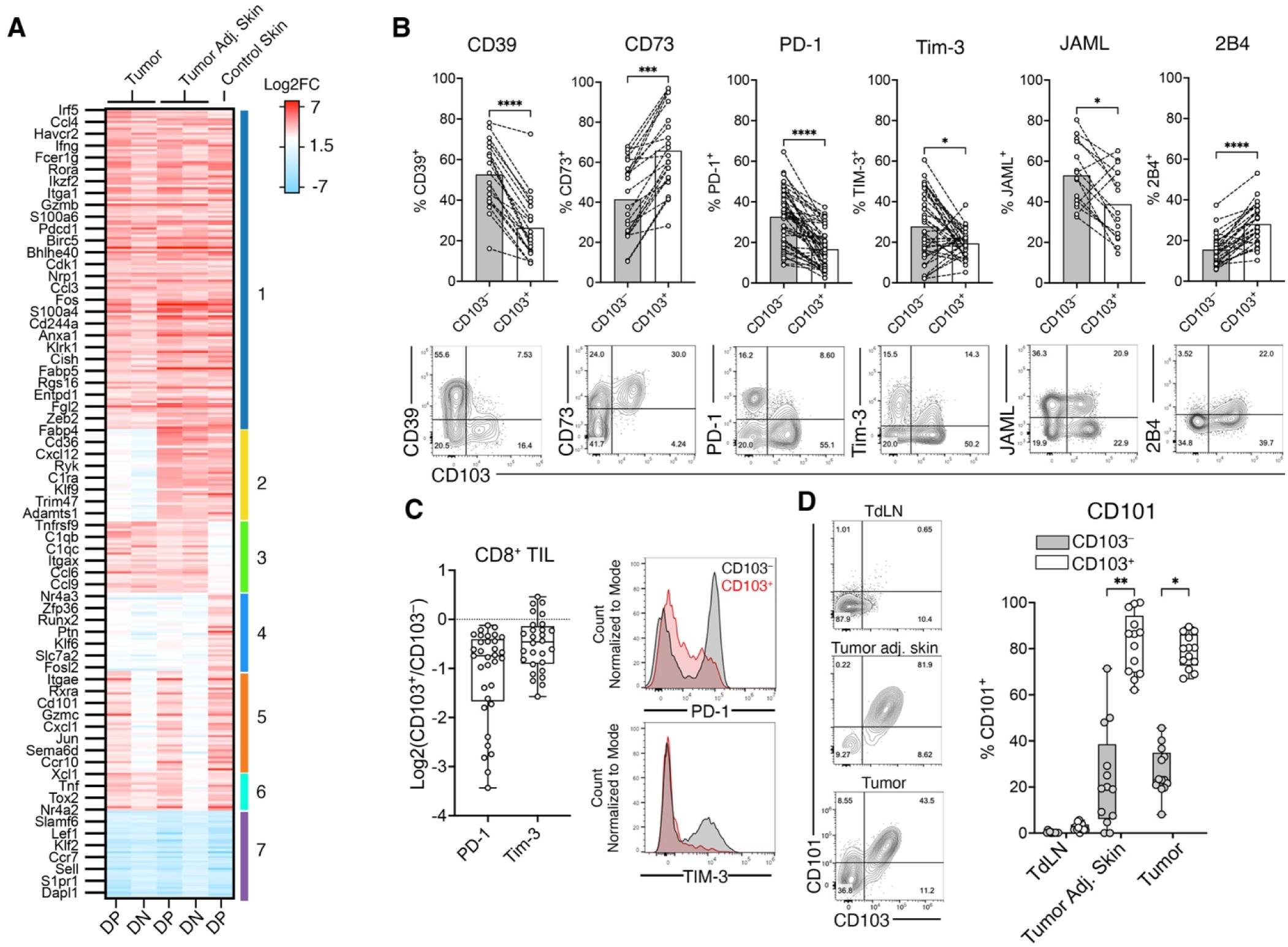
Phenotypic and transcriptional distinction of T_RM_-like versus CD103^−^ CD8^+^ TILs. **A**, Heatmap of differentially expressed genes (relative to splenic CD8^+^ T cells) from bulk RNA-Seq of DP (CD103^+^CD69^+^), DN (CD103^−^CD69^−^) CD8^+^ TILs and bona-fide skin T_RM_ controls. **B**, Flow cytometry of ecto-enzymes and co-inhibitory receptor expression by CD103^+^ and CD103^−^ CD8^+^ TILs. **C**, Comparison of PD-1 and Tim-3 median fluorescence intensity (MFI) in CD103^+^ versus CD103^−^ CD8^+^ TILs. **D**, Co-expression of CD101 with CD103 on CD8^+^ T cells from the tumor, tumor adjacent skin, and TdLN compartments. **B-D** display all mice from 2-5 independent experiments. Significance was determined by a Wilcoxon matched-pair signed rank test in **B** and **D.**

We then confirmed the level of protein expression for several of the cell surface receptors by flow cytometry. The ectoenzymes CD39 and CD73 displayed reciprocal expression patterns: CD39 was enriched in CD103^-^ cells, whereas CD73 was preferentially expressed in CD103^+^ T_RM_-like cells. Examination of coinhibitory receptors revealed that PD-1, Tim-3, and JAML were expressed at lower levels in T_RM_-like cells, while 2B4 was enriched in the CD103^+^ subset (**Figure 3B**), although the expression patterns of JAML and Tim-3 displayed variability across individual mice. Interestingly, PD-1 and Tim-3 on T_RM_-like TILs were expressed at intermediate levels, about 1.5-fold less based on median fluorescence intensity (MFI) compared to their CD103^−^ counterparts (**Figure 3C**). Strikingly, CD101, a type I transmembrane glycoprotein, was highly co-expressed with CD103 in both the tumor and tumor adjacent skin but not in the TdLN, suggesting CD101 as a potential marker to identify T_RM_-like cells (**Figure 3D**). Collectively, these findings define T_RM_-like CD8^+^ TILs as a phenotypically and transcriptionally distinct population within the tumor microenvironment.

### T_RM_-like cells compose the majority of the CD8^+^ TIL compartment in early tumors

We next investigated how the abundance of T_RM_-like CD8^+^ TILs changed with tumor progression. BPO tumors originate as multiple hyperpigmented lesions that increase in size and eventually coalesce into a single larger tumor. Histologically, these early lesions are in the radial growth phase (**Figure S3**). In some mice, both hyperpigmented lesions and a coalesced tumor can be found simultaneously, allowing for the analysis of multiple tumor stages within the same animal. We leveraged this observation to examine whether the stage of tumor development correlates with the presence of T_RM_-like CD8^+^ TILs. We compared tumor-adjacent skin (non-pigmented), hyperpigmented lesions (early-stage), and coalesced tumors (late-stage) from the same mouse. T_RM_-like cells constituted the majority of CD8^+^ TILs in hyperpigmented lesions, with frequencies comparable to those in tumor-adjacent skin (**Figure 4A**). We confirmed that these lesions contained transformed melanocytes by detecting EGFPOVA expression in the CD45^−^ fraction (**Figure 4B**). These findings suggest that CD8^+^ TILs are predominantly T_RM_-like during early tumor development.

**Fig. 4.**
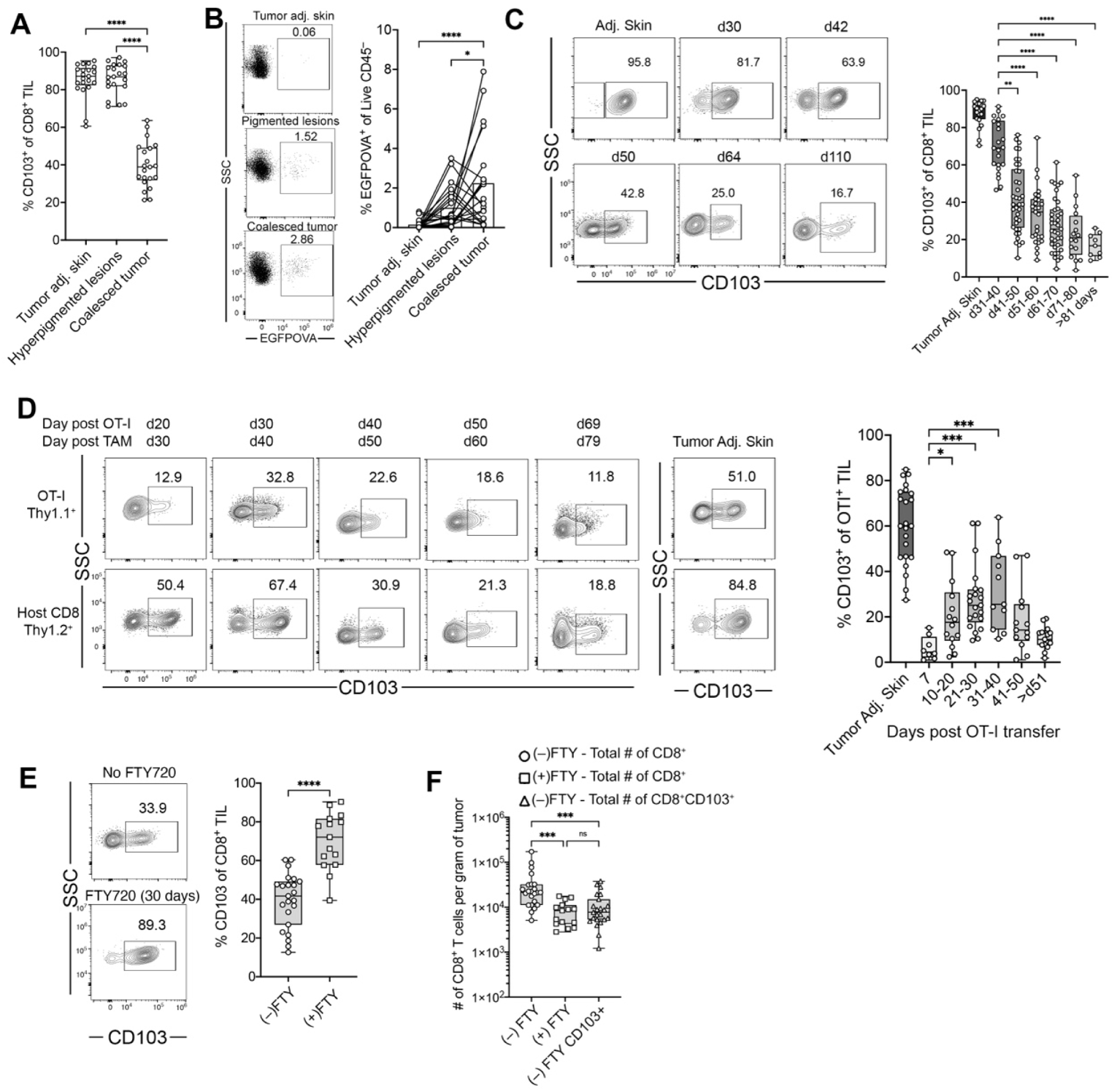
Dynamics of T_RM_-like CD8^+^ TILs during tumor progression. **A**, Frequencies of CD103^+^CD8^+^ TILs in tumor-adjacent skin, hyperpigmented lesions (early-stage) and coalesced (late-stage) tumors within the same mouse. **B**, EGFPOVA expression in CD45^−^cells from tumor adjacent skin, hyperpigmented lesions, and coalesced tumor within the same mouse. **C**, Time course of CD103^+^ CD8^+^ TIL frequencies binned by 10 day increments with representative flow cytometry plots. **D**, Expression of CD103 by OT-I and host CD8^+^ TILs over time. **E** and **F**, Representative flow plots showing the effect of FTY720 treatment on the relative frequency (**E**) and total number per gram of tumor of T_RM_-like CD8^+^ TILs (**F**) after 30 days. Graphs display all mice from 3 independent experiments. Significance was determined by a Kruskal-Wallis test with Dunn’s multiple comparison correction in **A-D** and **F** and a Mann-Whitney test in **E**.

To study this, we conducted a time-course analysis of T_RM_-like CD8^+^ TILs following tumor induction. In early-stage tumors (days 31–40 post-TAM), most CD8^+^ TILs expressed CD103 and CD101 (**Figure 4C** and **Figure S4A**). However, as tumors progressed, the frequency of T_RM_-like TILs steadily declined. By day 70, T_RM_-like TILs constituted only 20% of the CD8^+^ TIL compartment (**Figure 4C**). We extended this analysis to tumor-antigen-specific CD8^+^ TILs by adoptively transferring OT-I T cells at the onset of pigmentation. OT-I T cells upregulated CD103 after entering the tumor, peaking around day 30 post-transfer when approximately 30% expressed CD103. Similar to host (endogenous) polyclonal CD8^+^ TILs, OT-I cells expressing CD103 diminished in number over time, with only ∼12% expressing this marker in late-stage tumors (**Figure 4D**).

These data suggest that initial CD8^+^ T cell infiltrates preferentially acquire a T_RM_-like phenotype during early tumor development, but this population contracts during progression, due either to dilution by newly recruited non-T_RM_ cells or instability of the T_RM_-like state. To distinguish these possibilities, we sequestered T cells in tissues at the first signs of pigmentation by administering FTY720, a S1PR1 antagonist, in the drinking water; FTY720 prevents lymphocytes from trafficking out of lymphoid organs, preserving the tissue resident pool. We allowed the tumor to grow for 30 days and analyzed T cell frequencies and phenotypes. As expected, FTY720 reduced circulating T cells (**Figure S4B**) and decreased total CD8^+^ TIL numbers approximately tenfold. Intriguingly, the majority of CD8^+^ TILs were T_RM_-like in FTY720-treated mice (**Figure 4E**) and the absolute number of T_RM_-like CD8^+^ TILs remained comparable to untreated age-matched tumors (**Figures 4F**). These data suggest that, once established, the T_RM_-like population can persist locally for at least 30 days in the absence of continuous renewal from the circulation.

Taken together, these results suggest that the initial CD8^+^ T cell infiltrate into the tumor and tumor-adjacent skin differentiates into T_RM_-like cells, establishing a local niche. However, the gradual decline in antigen-specific T_RM_-like OT-I cells over time indicates a role for antigen-driven attrition as an additional component in determining T_RM_ abundance.

### Spatial analysis reveals early T_RM_-like cell activity in the epidermis and hair follicles and is associated with Treg cell infiltration

Given that T_RM_-like cells constitute the majority of CD8^+^ TILs in early tumors and express low levels of coinhibitory receptors, we hypothesized that alternative inhibitory mechanisms might be involved in regulating initial antitumor immune responses. To investigate this, we used t-CyCIF using an immune-focused murine antibody panel to determine immune organization and cellular interactions during early tumor evolution (**Figure S5A** and **S5B**). Due to substantial heterogeneity from one BPO mouse to the next, we biopsied the individual mice at three distinct stages: before pigmentation, at the first signs of pigmentation, and after tumor formation. To make this possible, tamoxifen was applied at two separate sites on the backs of mice, which served as the first two biopsy time points, and both sites were re-biopsied once tumors had fully developed (**Figure 5A**).

**Fig. 5.**
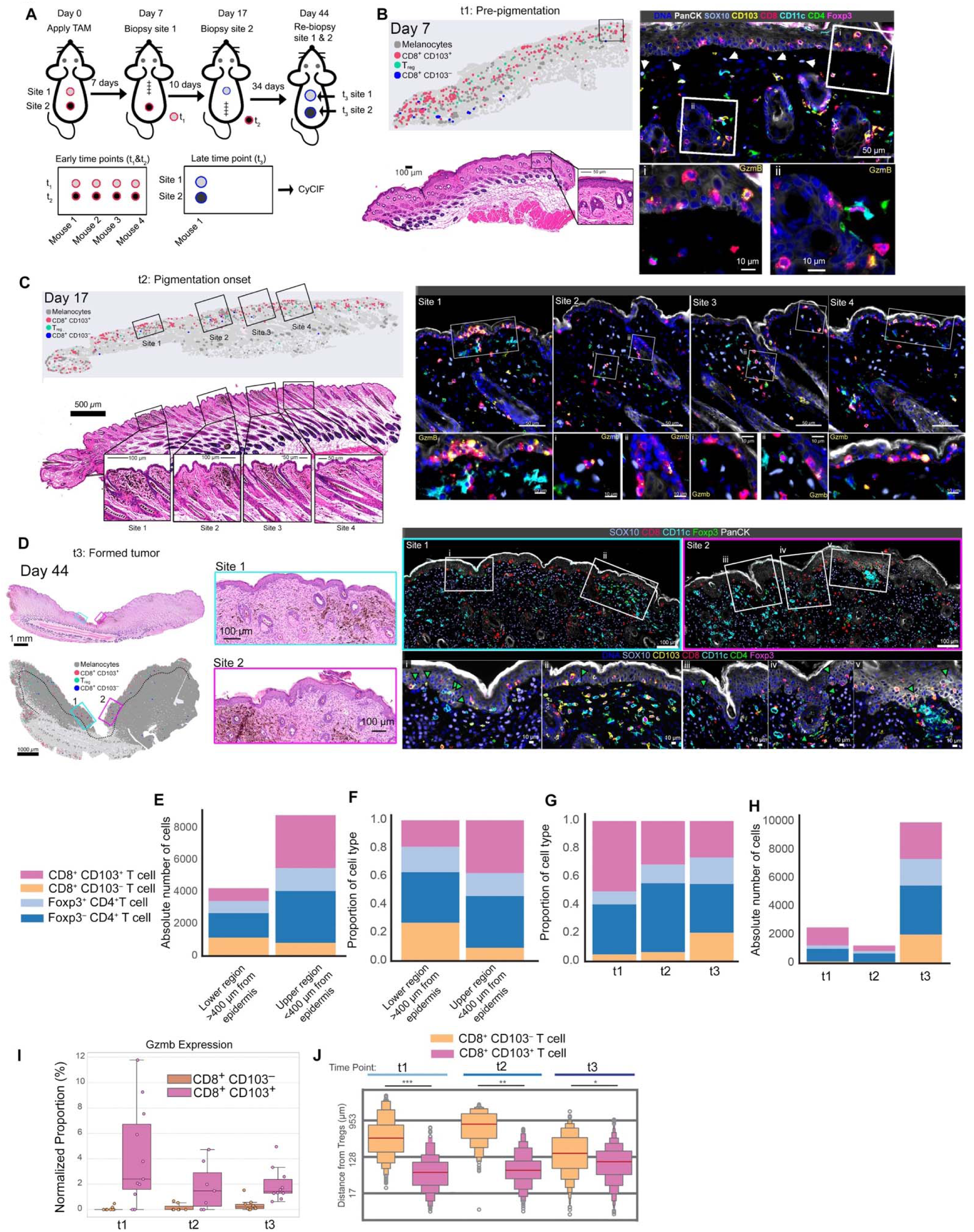
Spatial distribution and early activity of T_RM_-like cells and Treg infiltration by t-CyCIF. **A**, Experimental timeline and biopsy scheme (pre-pigmentation, pigmentation onset, and tumor formation). **B**, Representative scatter plot, multiplex immunofluorescence, and H&E image of a day 7 biopsy section. Insets highlight papillary dermal location of Sox10^+^ cells (white arrows), T_RM_-like and Treg cell localization, and GzmB expressed by T_RM_-like cells. **C**, Representative day 17 biopsy from the same mouse, with scatterplot, H&E stain, and multiplex imaging indicating hyperpigmented regions (sites 1-4). Insets show higher-magnification views of sites 1–4 highlighting epidermal location of CD8^+^ T_RM_-like cells and corresponding Gzmb expression. **D**, Scatterplot, H&E-stained sections, and multiplexed immunofluorescence of day 44 re-biopsy section from the same mouse. Insets (i-v) highlight Treg cell epidermal localization (green arrows) in proximity to CD8^+^ T_RM_-like cells. **E** and **F,** Absolute cell numbers (**E**) and relative proportions (**F**) of CD8^+^CD103^−^, CD8^+^CD103^+^, CD4^+^Foxp3^−^, and CD4^+^Foxp3^+^ T cells in the upper versus lower regions of the skin calculated among all time points and mice. **G** and **H,** Composition of CD8^+^CD103^−^, CD8^+^CD103^+^, CD4^+^Foxp3^−^, and CD4^+^Foxp3^+^ T cells across time points t1, t2, and t3, represented as relative proportions (**G**) and absolute cell numbers (**H**). **I**, Proportion of GzmB-expressing cells within CD8^+^CD103^+^ and CD8^+^CD103^−^ subsets across time points. **J**, Distribution of distances from Treg cells to CD8^+^CD103^+^ and CD8^+^CD103^−^ cells across time points, shown as violin plots with median and quartiles. Significance was determined by a Mann-Whitney U rank test in **J**.

At early time point (t1), Sox10^+^ cells were primarily localized within the lower hair follicle bulb, consistent with the presence of melanoblasts (**Figure S6A**), while scattered Sox10^+^ cells in the dermis and papillary dermis likely represented the earliest melanoma cells (**Figure 5B**, white arrows; **Figure S6B**). T cells and DCs were generally clustered around hair follicles and sparsely distributed in extra-follicular space (**Figure 5B**). The epidermis primarily contained T_RM_-like CD8^+^ T cells, which often expressed GzmB (**Figure 5B** and **5I**).

At the second time point (t2), we observed a slight increase in the frequency of Sox10^+^ cells overall (**Figure S5C** and **S5D**), and the appearance of clusters of Sox10^+^ melanoma cells, which often colocalized with hyperpigmented regions (**Figure 5C** and **Figure S7**), but could also be amelanotic (**Figure S7B**, site iv). In general, these Sox10^+^ melanoma cell clusters were enriched for immune cells including GzmB^+^ T_RM_-like cells, Treg cells, and DCs (**Figure 5C**, sites 1-4, and **S7**; sites i-iii and v) but varied in the composition and spatial distribution of immune cells, even within the same mouse. For example, some clusters contained higher densities of CD4^+^Foxp3^+^ Treg cells compared to CD8^+^ TILs (**Figure S7B**, site iv and **S7C**, site vi) or were macrophage-dominant with few CD8^+^ T cells (**Figure S7**, site vii).

In late-stage tumors (t3), some regions remained pigmented, however, the bulk of the invading mass exhibited loss of pigmentation (**Figure 5D**). Sox10^+^ melanoma cells had invaded deep into the hypodermis, and nearly all mice contained LN metastases (data not shown). Interestingly, most T cells appeared to reside in the upper regions of the skin and were only sparsely distributed deeper in the tumor bed (**Figure 5D**): indeed, quantification of CD8^+^ (CD103^+^ and CD103^−^) T cells across all time points revealed approximately 2-fold more T cells within ∼400 µm of the epidermis, with T_RM_-like cells abundant in the upper regions (**Figure 5D-F**). Consistent with our flow cytometry data, T_RM_-like CD8^+^ T cells were most abundant at early stages, but their frequency diminished as the tumor progressed (**Figure 5G**). Additionally, between the early (t1 and t2) and late (t3) time points, we observed an increase in the total number of CD8^+^ CD103^−^, Tconv, and Treg cells, while T_RM_-like cells remained constant in number, which supports the notion that distinct T_RM_-niches exist within the TME and are filled early in immune cell recruitment (**Figure 5H**). Analysis of GzmB expression revealed that T_RM_-like cells comprised the majority of GzmB^+^ CD8^+^ T cells across all time points but their relative frequency diminished as the tumor progressed (**Figure 5I**). Finally, we found that at all time points, T_RM_-like CD8^+^ TILs were in closer proximity to Treg cells as compared to their CD103^−^counterparts (**Figure 5J**). However, in late-stage tumors CD103^−^ CD8^+^ TILs were closer to Treg cells compared to earlier time points, likely reflecting the overall increase in Treg cell numbers. Additionally, in late-stage tumors, Treg cells could be found within the epidermis, which was rare in earlier time points (**Figure 5D**, sites i-v, green arrows), indicating an activation of Treg cell migration, which can occur during inflammation.

Taken together, these data suggest that T_RM_-like CD8^+^ T cells are highly active in the epidermis and hair follicles during early melanoma development, where they are closely associated with infiltrating regulatory T cells, suggesting early immune crosstalk and niche formation within the tumor microenvironment.

### T_RM_-like cells immunoedit tumors early in the antitumor immune response

Since Gzmb was upregulated in T_RM_-like cells within hyperpigmented lesions, we hypothesized that these cells might play a role in immunoediting during the earliest stages of tumor development. To investigate this, we first assessed whether T_RM_-like cells were tumor-specific using H2-K^b^/SIIN-pentamer analysis at both early and late time points. Unexpectedly, while a subset of T_RM_-like cells were H2-K^b^/SIIN-specific, the CD103^−^ population contained a higher frequency of pentamer binding cells, which was further enriched in late tumors (**Figure S8A**). This pattern reflects the observed decrease in CD103 expression over time in the OT-I transfer study (**Figure 4D**) and suggests instability within the T_RM_-like population, possibly driven by antigen stimulation. To determine whether T cells present in the tumor during early development, which are predominantly T_RM_-like cells, contribute to immunoediting, we depleted CD8^+^ T cells using an anti-CD8α antibody beginning at the first signs of pigmentation and continuing every five days until the end of the experiment (**Figure S8B**). We confirmed a reduction in both the frequency and total number of CD8^+^ TILs at the endpoint (**Figure S8C**). Using EGFPOVA as a marker for non-immunoedited tumor cells, we found that tumors in CD8-depleted mice contained a higher frequency of EGFPOVA^+^ tumor cells (**Figure S8D**). This frequency was comparable to tumors from immunodeficient BPO/Rag1^−/–^ mice. Thus, by reducing the frequency of T_RM_-like cells early in the antitumor immune response we blunted CD8 T cell-mediated immunoediting, suggesting that T_RM_-like cells contribute to early tumor immunoediting.

### Treg depletion enhances tumor control and expands tumor-specific CD8^+^ T Cells

Given the presence of Treg cells and their co-localization with T_RM_-like cells early in tumor development, we asked whether Treg depletion would enhance tumor control potentially by activating the T_RM_-like population. To do this, we crossed BPO mice with Foxp3DTR-EGFP mice; in these animals, which express human diphtheria toxin receptor, exposure to diphtheria toxin (DT) results in ablation of Foxp3^+^ Treg cells. We first assessed Treg cell expression of the activation markers PD-1, Tim-3, and CTLA-4 in the TdLN, tumor-adjacent skin, early tumors, and late tumors. While these markers were expressed at low levels in Tregs in the TdLN, their expression progressively increased from tumor-adjacent skin to early tumors and reached the highest levels in late tumors (**Figure 6A**). Most Treg cells expressed CD103 in the tumor-adjacent skin, early tumors, and late tumors, further supporting their ability to co-localize with T_RM_-like CD8^+^ TILs (**Figure 6A**). In parallel with increasing Treg expression of activation markers, we also observed a progressive increase in Treg cell abundance across these sites (**Figure 6B**). As a consequence, CD8^+^ T cells were more abundant in the tumor-adjacent skin but in early tumors Treg and CD8 T cells were found at similar frequencies and late-stage tumors contained more Treg cells than CD8 T cells (**Figure 6C**).

**Fig. 6.**
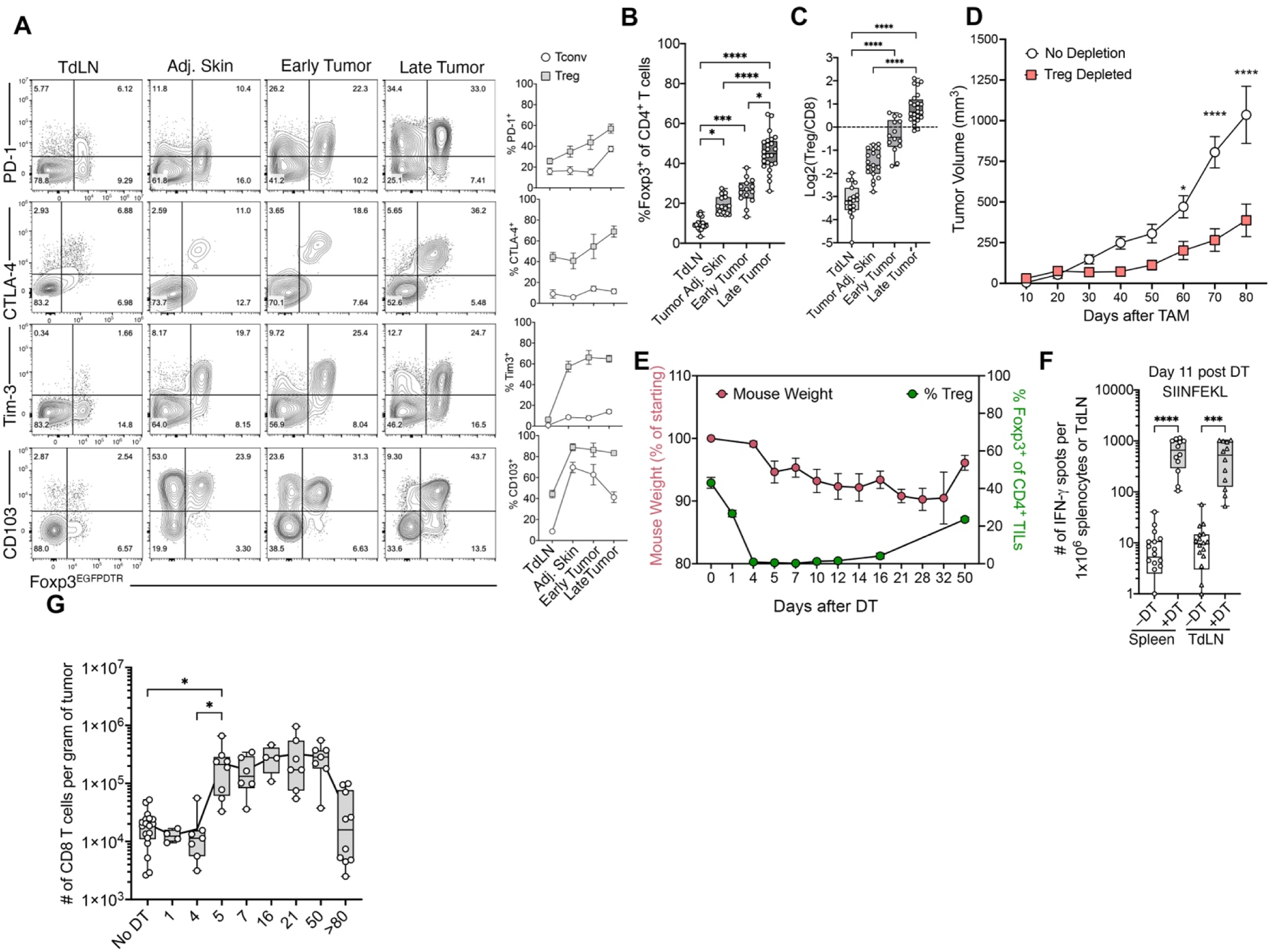
Systemic Treg depletion enhances tumor control and expands tumor-specific CD8^+^ T cells. **A**, PD-1, Tim-3, CTLA-4 and CD103 expression on Treg cells in TdLN, tumor adjacent skin, early and late tumors. **B** and **C**, Treg cell abundance (**B**) and Treg/CD8 ratio (**C**) in TdLN, tumor adjacent skin, early and late tumors. **D**, Tumor growth curves following two DT doses (i.p.) versus control. **E**, Treg cell kinetics in the tumor and mouse weight after DT treatment. **F**, IFN-γ ELISPOT of anti-SIINFEKL T cell response on day 11 post-DT administration. **G**, Total number of CD8 TILs cells following Treg depletion. Graphs display all mice from 2-5 independent experiments. Significance was determined by a Kruskal-Wallis test with Dunn’s multiple comparison correction in **B**, **C**, **F**, and **G**. For tumor outgrowth (**D**) a Two-way ANOVA with Sidak multiple comparisons correction was used.

To determine whether transient Treg depletion would induce tumor control and enhance the antitumor immune response we administered two doses of DT i.p. into tumor bearing mice and monitored tumor growth. We observed that Tregs were nearly absent in the tumor by day 3 post-DT treatment and remained depleted for 14 days. In these animals, tumor growth was significantly reduced (**Figure 6D**). Treg numbers rebounded by day 39 but their frequency never returned to pre-depletion levels (**Figure 6E**). As expected, Treg cell-depleted mice experienced systemic autoimmunity highlighted by rapid weight loss (**Figure 6E**), but most mice eventually recovered. Strikingly, analysis of the antitumor immune response directed against the OVA epitope SIINFEKL 11 days post-depletion revealed a ∼100-fold increase in the frequency of tumor-specific CD8^+^ T cells in both the spleen and TdLN (**Figure 6F**). Tracking the frequency of CD8^+^ T cells in the tumor following Treg depletion showed a ∼10-fold increase in the total number of CD8^+^ TILs by day 5, and they remained elevated for at least 50 days (**Figure 6G**). We also probed for changes in other cellular compartments in the tumor including CD4 T cells, γδ T cells, DCs, macrophages, NK cells, and Neutrophils. Other than CD8 T cells, only CD4 T cells exhibited a significant increase in frequency (**Figure S9A**). These data show that depletion of Treg cells remodels the tumor microenvironment, slowing tumor growth while driving robust expansion of tumor-specific CD8^+^ T cells in the periphery and enhanced infiltration into the tumor.

### Tumor site-specific depletion of Treg cells activates T_RM_-like cells

To determine whether the increase in CD8^+^ tumor-infiltrating lymphocytes (TILs) resulted from local proliferation or tumor infiltration, we depleted Tregs followed by daily FTY720 administration to block T cell trafficking, and quantified CD8^+^ TILs 5 days later. The increase in CD8^+^ TILs caused by Treg depletion was completely abrogated in FTY720-treated mice, showing that the increase in TILs was a result of CD8^+^ T cell recruitment (**Figure 7A**). A similar effect was observed for CD4^+^ Foxp3^−^ T cells (**Figure S9B**). We also observed upregulation of PD-1 in both the CD8^+^ CD103^−^ and CD103^+^ subpopulations (**Figure 7B**); a similar change was observed for Tim-3 expression (**Figure S9C**). Although bulk PD-1 upregulation likely reflects both tumor-residing and newly recruited CD8^+^ T cells, the retention of PD-1 upregulation under FTY720 demonstrates that tumor-residing CD8^+^ T cells contribute directly to this response.

**Fig. 7.**
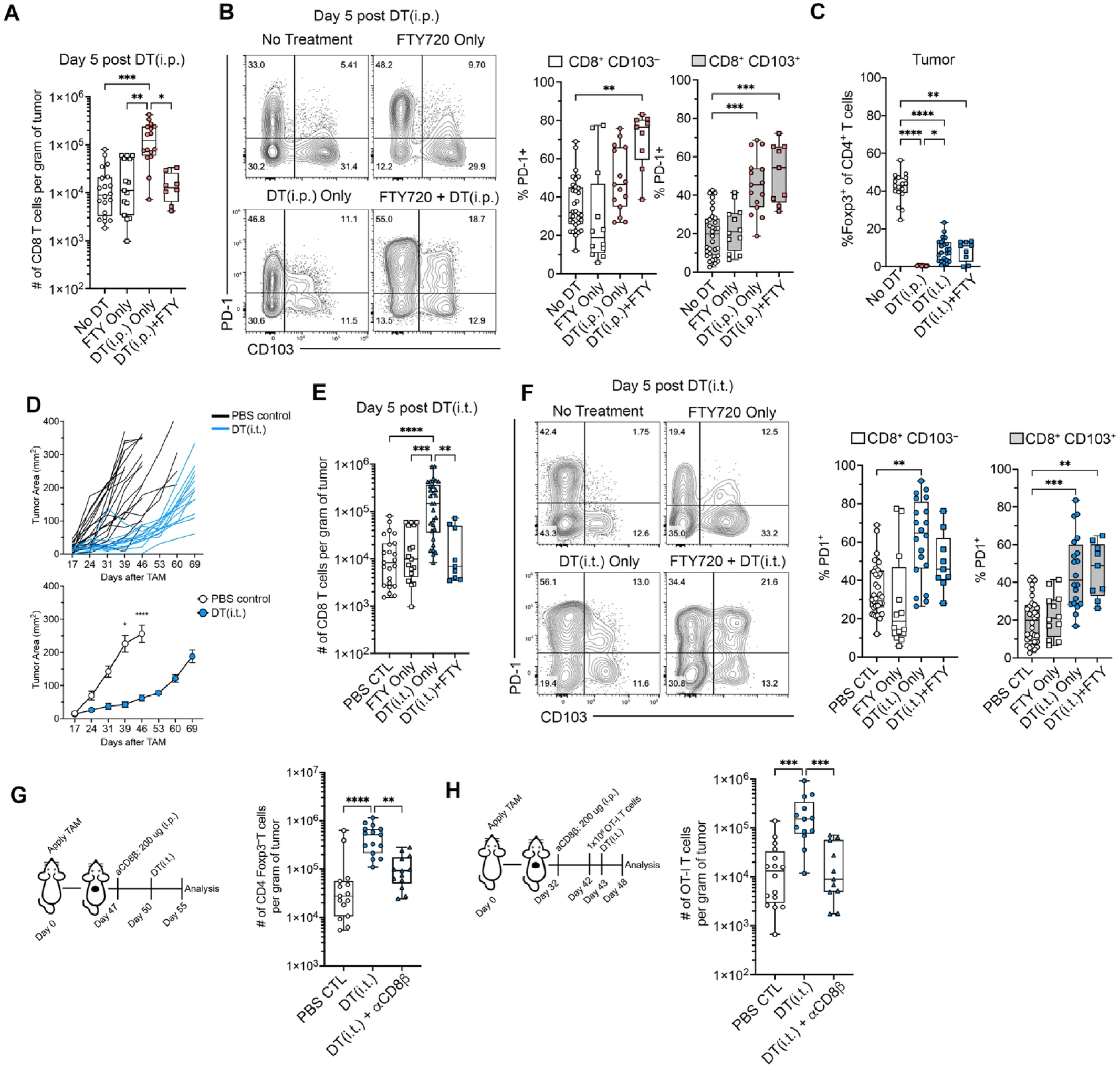
Local Treg depletion activates T_RM_-like cells and promotes T cell recruitment. **A**, Number of CD8^+^ TILs per gram of tumor on day 5 following systemic Treg depletion (DT i.p.) ± FTY720. **B**, Representative flow cytometry plots and quantification of PD-1 expression on CD8^+^ CD103^−^ and CD8^+^ CD103^+^ TIL subsets on day 5 following DT(i.p.) ± FTY720. **C**, Frequency of Foxp3^+^ cells among tumor CD4^+^ T cells following systemic (i.p.) or intratumoral (i.t.) DT administration ± FTY720. **D**, Tumor area over time in mice receiving weekly intratumoral DT (DT(i.t.), 12.5 µg/kg, once per week) versus PBS-treated controls, shown as individual mouse growth curves (top) and group means ± SEM (bottom). **E**, Number of CD8^+^ TILs per gram of tumor on day 5 following intratumoral Treg depletion (DT i.t.) ± FTY720. **F**, Representative flow cytometry plots and quantification of PD-1 expression on CD8^+^ CD103^−^ and CD8^+^ CD103^+^ TIL subsets on day 5 following DT(i.t.) ± FTY720. **G**, Experimental schematic (left) and quantification (right) of CD4^+^ Foxp3^−^ Tconv cell recruitment into the tumor following DT(i.t.)-mediated Treg depletion, with or without prior systemic CD8^+^ T cell depletion using anti-CD8β antibody (200 µg, i.p.). **H**, Experimental schematic (left) and quantification (right) of OT-I T cell recruitment into the tumor following adoptive transfer of 1x10^6^ OT-I T cells and DT(i.t.)-mediated Treg depletion, with or without prior systemic CD8^+^ T cell depletion. Graphs display all mice from 2–5 independent experiments. Significance was determined by a Kruskal-Wallis test with Dunn’s multiple comparison correction in **A**, **B**, **C**, **E**, **F**, **G**, and **H**. For tumor outgrowth (**D**), a Two-way ANOVA with Sidak’s multiple comparisons correction was used.

A limitation of systemic Treg depletion is the potential for widespread autoimmunity, and the release of cytokines into the circulation which may non-specifically influence tumor-residing T cells (*33*, *34*). To achieve local Treg depletion, while minimizing systemic autoimmunity, we administered a lower dose of DT (12.5 µg/kg) intratumorally once every seven days, which did not cause weight loss (**Figure S9D**) or eliminate Treg cells in the TdLN (**Figure S9E**). However, we observed an approximately fivefold reduction in Treg cell frequency within the tumor, though depletion was less complete as compared to DT administered intraperitoneally (**Figure 7C**). Nonetheless, weekly intratumoral DT administration significantly restrained tumor outgrowth as compared to PBS-treated controls (**Figure 7D and S9F**). Serial digital photography revealed eradication of early hyperpigmented lesions in DT(i.t.)-treated mice, which was frequently accompanied by the development of vitiligo at the treatment site (**Figure S10**).

Low dose DT(i.t.) provided a system to address two questions: whether tumor-restricted Treg depletion, leaving peripheral Treg cells intact, was sufficient to drive CD8^+^ T cell recruitment and systemic antitumor immunity, and whether the incomplete intratumoral Treg loss achieved by this regimen could still reactivate T_RM_-like cells. To this end, we combined DT(i.t.) with FTY720 administration. FTY720 did not alter the extent of intratumoral Treg depletion in the tumor (**Figure 7C**) and TdLN (**Figure S9E**). DT(i.t.) drove significant CD8^+^ and CD4+ Foxp3^−^ T cell recruitment into the tumor, which was blocked by FTY720 (**Figure 7E** and **S9F**). T_RM_-like CD8 TILs still exhibited upregulation of PD-1 (**Figure 7F**) and Tim-3 (**Figure S9H**), further demonstrating that the change in CD8 T cell state was a direct effect of local Treg depletion. In parallel, DT(i.t.) drove a significant expansion of SIINFEKL-reactive CD8^+^ T cells in the spleen and the TdLN as measured by IFN-γ ELISPOT (**Figure S9G**).

The increase in PD-1 and Tim-3 expression after Treg depletion suggested that T_RM_-like CD8^+^ TILs had become activated. In infection models, a key sentinel function of T_RM_ cells is the recruitment of other immune cells to inflamed tissues (*12*, *14*), a process thought to be mediated by IFN-γ release. Additionally, conventional type 1 dendritic cells (cDC1s) have been shown to promote T cell recruitment to the tumor microenvironment (*35*). To investigate whether CD8^+^ T cells contribute to immune cell recruitment, we depleted CD8^+^ T cells with systemic (i.p.) administration of an anti-CD8β antibody to avoid depleting cDC1s, which can express CD8α. 72 hours later, we depleted Treg cells with DT(i.t.), and 5 days post-Treg depletion, we quantified CD4^+^ Foxp3^−^ Tconv cell infiltration as a measure of T cell recruitment, while also confirming a reduction in total CD8^+^ TIL numbers at this endpoint (**Figure S9I**). Surprisingly, CD8^+^ T cell depletion blunted the recruitment effect of Tconv cells following Treg depletion (**Figure 7G**). To test if tumor-residing CD8^+^ T cells were involved in peripheral CD8 T cell recruitment, we waited 10 days for free anti-CD8β antibody levels to diminish before transferring 1 x 10^6^ OT-I T cells into tumor-bearing BPO/Foxp3DTR mice followed by DT(i.t.)-mediated Treg depletion. As in the Tconv experiment, we confirmed reduced host CD8^+^ TIL numbers at endpoint (**Figure S9I**), and observed that OT-I CD8^+^ T cell recruitment into the tumor was diminished with prior host CD8 T cell depletion (**Figure 7H**). Lingering anti-CD8β antibody likely did not deplete infiltrating OT-I T cells as the number of OT-I T cells in the TdLN was not diminished (**Figure S9J**). Taken together, these data demonstrate that T_RM_-like CD8^+^ T cells, once released from Treg-mediated suppression, function as local sentinels: driving recruitment of additional effector T cells into the tumor.

## Discussion

T_RM_ cells are a unique lineage of T cells with specialized functions that endow them with the capacity to adapt, survive, and exert potent effector functions in their tissue of residence. T cells expressing markers of tissue-residency or containing a T_RM_ transcriptional profile are found across solid tumor types and their presence is associated with improved survival and response to immunotherapy (*36–38*). T_RM_-like T cells in tumors can be either pre-existing, for example, those that populate tissue in response to infection, or tumor reactive, those that are primed in the TdLN, infiltrate into the tumor, and differentiate into T_RM_-like cells. A subset of pre-existing CD8^+^ TILs is pathogen-specific (*39–41*), but whether the immune system generates T_RM_-cells in response to a progressing tumor is largely unknown. Preclinical studies of T_RM_ cells in cancer have, for the most part, involved prophylactic T_RM_ cell seeding before tumor challenge. While critical for uncovering the potent tumoricidal properties of T_RM_ cells, it is unclear whether an infiltrating CD8^+^ T cell enters the T_RM_ lineage. Here, we show that the earliest stages of tumor development involve T_RM_-like CD8^+^ T cells interacting with an initializing tumor microenvironment. We do not believe these T_RM_-like cells to be pre-existing as mice are kept in a pathogen-free environment, however, mouse skin is populated with a small number of T_RM_ cells after birth (*42*). Therefore, further investigation is needed to definitively determine the origin of these cells.

The T_RM_-like population was identified by expression of CD103. This population was phenotypically distinct from the CD103^−^ population and expressed lower levels of the coinhibitory receptors PD-1 and Tim-3, as well as differential expression of the ecto-nucleosides CD39 and CD73, and co-expression of CD101. A similar pattern of expression was recently reported in CD8^+^ T_RM_ cells in human breast cancer (*43*). Intriguingly, CD101 has been found on T_RM_ cells in multiple tissues (*10*, *44*, *45*) and recently reported to preferentially identify epidermal CD8^+^ T_RM_ with IFN-γ production capacity in human skin (*46*). While the function of CD101 *in vivo* is unknown, it was shown to suppress T cell function *in vitro* (*47*) and was identified in terminally exhausted CD8 T cells in a model of chronical viral infection (*48*). Therefore, whether CD101 is a targetable receptor to activate T_RM_-like cells in tumors warrants further investigation.

In human tumors, T_RM_-like cells were found to express CD39 and PD-1 (*49*). One explanation as to why T_RM_-like CD8^+^ TILs in the BPO model do not follow this pattern may be the relative activity of Treg cells, as PD-1 expression was upregulated in mice when Treg cells were depleted. Alternatively, while tumor-specific SIIN/H-2Kb^+^ CD8^+^ TILs were found in both CD103^+^ and CD103^−^ populations at early time points, the majority of SIIN/H-2Kb^+^ CD8^+^ TILs were CD103^−^ at later time points. This, coupled with our observation that EGFPOVA^+^ tumor cells are rapidly eliminated in most tumors, suggests either that antigen-availability promotes the T_RM_-like population or high-affinity TCR stimulation is unfavorable to generate T_RM_-like CD8^+^ TILs. Fully understanding the relationship between the CD103^+^ and CD103^−^ population will require lineage tracing studies.

We found that most T_RM_-like CD8^+^ T cells remained in and near the epidermis throughout tumor progression, while a sporadic few were found deeper in the tumor bed. This suggests that signals driving this population, possibly TGFβ (*17*, *50*), are expressed in the upper regions of the skin. It also suggests that a niche for T_RM_-like cells exists in tumors. In support of this idea, preventing new T cell recruitment with FTY720, starting when hyperpigmented lesions appear, showed that the T_RM_-like population can persist in a progressing tumor for at least 30 days. After 30 days, the total number of T cells is similar to the total number of T_RM_-like CD8^+^ TILs in untreated mice.

Depletion of CD8 T cells at the time of pigmentation increased the frequency of EGFPOVA^+^ tumor cells, suggesting tumoricidal activity in the T_RM_-like population. Furthermore, T_RM_-like CD8^+^ TILs upregulated Gzmb during the early stages of tumor development, suggesting cytolytic activity. However, the increase in EGFPOVA^+^ tumor cells was modest. Yet, it did match the frequency of EGFPOVA^+^ tumor cells from BPO/Rag1^−/–^ mice. To explain the relatively low baseline EGFPOVA expression and the modest increase after CD8 T cell depletion, we propose two possibilities; first, it is possible that Cre activity after 4-OHT application does not lead to EGFPOVA expression in 100% of transformed cells, and thus only a fraction of tumor cells express EGFPOVA. Second, it’s known that bona fide T_RM_ cells in the skin at homeostasis are not effectively depleted with antibody. Therefore, it is possible that our depletion was incomplete, and the remaining T_RM_-like cells were able to eliminate tumor cells. In addition, non-T_RM_-like CD8^+^ TILs likely contribute to immunoediting and may be the dominant cytotoxic CD8 T cell population at later time points after the tumor escapes the initial T_RM_-mediated T cell insult.

It is known that tumors in the Braf/PTEN melanoma model contain a relatively low level of T cell infiltrate, and the T cell infiltrate is skewed towards CD4 and Treg cells. Interestingly, BPO-derived tumor cell lines engrafted subcutaneously develop tumors that generally contain a greater CD8/CD4 T cell ratio (data not shown), compared to autochthonous BPO tumors. Part of this is likely explained by an immunogenic start to subcutaneous engrafted tumors driven by large degree of tumor cell death after injection. Also, it suggests that the natural development of melanomas in the autochthonous setting influences the magnitude of the T cell infiltrate. In the absence of an immunogenic tumor initiation, our data suggest that Treg cells play a critical role in dampening early CD8^+^ T cell recruitment to the tumor. The source of Treg cells can also be either pre-existing or recruited. It is known that oncogenic Braf can drive CCR4-dependent recruitment of Treg cells to the tumor (*51*). In agreement, we observed a roughly 2-fold increase in Treg cell abundance between early and late tumors. Additionally, after birth, Treg cells localize to hair follicles and promote immune tolerance where hair follicle stem cells reside. Therefore, pre-existing Treg cells may also play a role in dampening the initial antitumor T cell response (*52–54*).

A central question raised by our findings is whether reactivation of the early T_RM_-like infiltrate is sufficient to alter the course of tumor growth. Strikingly, tumor-restricted Treg depletion was enough to restrain outgrowth: weekly intratumoral low-dose DT, which left peripheral Treg cells intact and achieved only partial intratumoral Treg loss, significantly slowed tumor growth and, in many animals, eradicated early hyperpigmented lesions outright. The frequent appearance of vitiligo at treated sites further indicates that releasing the early infiltrate from Treg restraint generates productive, durable antitumor immunity. The result is a remodeling of the T cell infiltrate from a CD4/Treg-dominated state to one that recruits and amplifies effector CD8 T cell responses, positioning the early T_RM_-like:Treg axis as an initial barrier to antitumor immunity, imposed at tumor onset and well before exhaustion programs take hold. Perturbing this axis locally augments CD8 T cell immunity systemically, making it an attractive immunotherapeutic target.

Our data support the notion that part of the Treg-mediated inhibition of the antitumor T cell response operates through the silencing of T_RM_ “tissue alarm” functions. In infection models, the T_RM_ alarm function is mediated through IFN-γ, which acts on nearby cells to secrete T cell recruiting chemokines. Future studies will determine whether T_RM_-secreted IFN-γ is suppressed by Treg cells and if activating T_RM_-like cells can induce T cell recruitment in the presence of Treg cells. We propose that the initial T cell response to melanoma follows a tissue residency trajectory with CD8 T cells entering a T_RM_-like state, exhibiting cytotoxic and alarm functions. This T_RM_ response is rapidly suppressed by Treg cells but can resume when Treg cells are eliminated.

## Method Details

### Mice

B6.Cg-Tg(Tyr-cre/ERT2)13Bos *Braf^tm1Mmcm^ Pten^tm1Hwu^*/BosJ mice were purchased from Jackson and crossed to an inducible Rosa26-GFPOVA (*31*), a gift from Dr. Angelika Sales (formerly Stoecklinger) at the University of Salzburg, to generate Braf^CA/WT^/PTEN^loxp^/R26-EGFPOVA^+/+^/TyrCreERT2 (abbreviated BPO) mice. BPO mice were crossed to Foxp3DTR mice (B6.129(Cg)-*Foxp3^tm3(Hbegf/GFP)Ayr^*/J) to generate BPO/Foxp3DTR mice. All BPO mice used were heterozygous for BrafV600E (Braf^CA/WT^). OT-I/Rag1^−/–^ /Thy1.1 mice were maintained in house. For subcutaneously engrafted tumor cell lines BPO TyrCre-negative mice were used. Both male and female mice were used. All animal work and protocols in the current study were approved by the IACUC committee at the Center for Comparative Medicine at Brigham and Women’s Hospital, accredited by AAALAC.

### Clinical samples

All CyCIF images with associated histopathological annotations for Stage II melanoma cohort were processed and imaged as described in Vallius et al. 2025 (doi: https://doi.org/10.1101/2025.06.21.660851). Full-resolution CyCIF images, single cell segmentation masks, and cell count tables are available via the NCI Human Tumor Atlas Network data portal (https://data.humantumoratlas.org/). Code used for multimodal spatial analysis are available on GitHub (https://github.com/labsyspharm/2025_Williams_Pant_Melanoma-TRM). Based on the melanoma diagnostic criteria, the histopathologic annotations included normal skin (N), Precursor (P), and Vertical Growth Phase melanoma (VGP).

### Cellular depletions, FTY720 treatment, adoptive cell transfers

CD8 T cells were depleted by administration of 200 ug/dose of anti-CD8α (clone 2.43, BioXCell) or anti-CD8β (53-5.8, BioXCell) antibody. For systemic Treg cell depletion, 50 μg/kg DT was administered i.p. For intratumoral Treg depletion, mice were anesthetized, and tumors were injected with 12.5 μg/kg DT intratumorally in a total volume of 50 ul. FTY720 was administered by either daily i.p. injection of 1.25 mg/kg or by addition to the drinking water to a final concentration of 3 ug/mL. OT-I T cells were adoptively transferred retro-orbitally. For proliferation OT-I cells were labeled with Cell Trace Violet according to manufacturer’s protocol.

### Autochthonous tumor induction, tissue harvest, and cell line generation

To induce tumors, a 1 cm x 1 cm area on the backs of mice was shaved and depilated using Veet cream. The area was washed with ethanol and 1 ul of 4-hydroxytamoxifen (dissolved in DMSO to a concentration of 100 mg/mL). Tumors were induced at 6-9 weeks after birth. Mice were monitored weekly for tumor formation. If an off-site non-4-OHT-induced tumor was found at the start of the experiment the mouse was removed from the study. For systemic Treg depletion tumor length (T_L_), width (T_W_), and height (T_H_) were measured with digital caliper and tumor volume was calculated (T_vol_ = T_L_ x T_W_ x T_H_). For DT(i.t.) experiments, a handheld digital microscope (DinoLite) was used to image tumor lesions and capture changes in tumor morphology over time; tumor area was calculated from these images (Dino Capture 2.0). Tissue was harvested by depilating the hair on the backs of mice with Veet and washing the area with 70% ethanol. Tumor, tumor adjacent skin, and hyperpigmented lesions were excised with scissors and placed in a 70 mm tissue culture dish. Tumor adjacent skin and hyperpigmented lesions were minced with scissors and transferred to a 5 mL tube containing 1 mL digestion buffer (collagenase, type IV (10 mg/mL), hyaluronidase (1 mg/mL), and DNase (200 mg/mL) in HBSS). Tumors were injected with 1 mL digestion buffer, minced with scissors, and transferred to a 5 mL tube. Samples were incubated at 37°C with rotation for 30 min then transferred to a 100 um filter and systematically pushed through the filter using a syringe plunger with 4 filter washes using 5 mL of tumor wash buffer (PBS with 1% FBS, 1 mM EDTA, and 1X penicillin/streptomycin). Samples were spun and refiltered through a 100 um filter into a 15 mL conical tube. Live cells were purified by Ficoll and washed twice before antibody staining. To generate tumor cell lines, the pellet from the Ficoll step was filtered through a 100 um mesh filter and cultured in vitro. Approximately 16% of tumor samples generated a tumor cell line that grew progressively in vitro and in vivo.

### BPO-derived tumor cell lines and engraftment

All cell lines were grown in DMEM supplemented with 10% fetal bovine serum, MOPS (10 mM), and Penicillin/Streptomycin (100 U/mL) under 37°C / 5% CO2 conditions. BPO-derived tumor cell lines were generated in-house by extended in vitro cell culture without in vivo passaging in immune compromised mice. Cell lines were generated from male and female mice. B16.OVA and MC38.OVA cells were generated by lentiviral transduction of pLV-CMV>{OVA}:3xGGGGS:mCherry (Vector Builder) and sorted to 100% purity based on mCherry expression. Tumor cells were engrafted on the flanks of mice and tumor volume (T_L_ x T_W_ x T_H_) was measured with a digital caliper.

### Bulk RNASeq

Library preparation and RNA sequencing was performed by the Dana-Farber Molecular Biology Core Facilities using Takara SmartSeq v4 reagents for low input mRNASeq. Samples with an RNA quality (RIN) score >7 were used for sequencing. Full-length cDNA was fragmented to an average size of 200 bp, and sequencing libraries were prepared from 2 ng of sheared cDNA. Double-stranded DNA libraries were quantified by Qubit fluorometer and Agilent TapeStation 2200. Uniquely dual-indexed libraries were pooled in equimolar amounts, assessed for cluster efficiency and pool balance with shallow sequencing on an Illumina MiSeq. Final sequencing was performed on an Illumina NovaSeq X Plus with paired-end 150bp reads. Sequencing data was processed and analyzed using Partek software. Alignment was performed by the STAR (version 2.7.8a) alignment method to mouse Genome assembly GRCm39. Raw read counts were processed by TMM normalization. Differentially expressed genes were determined by normalization to spleen CD8 samples with a cutoff fold-change > 2 and adjusted p-value <0.05.

### Flow cytometry data acquisition and analysis

For bulk RNASeq, BPO tumors, tumor adjacent skin, and spleens from 3 BPO mice were pooled and sorted for CD8 T cells expressing CD103 and CD69. For the bona fide T_RM_ control, Yumm1.7.OVA cells were engrafted epicutaneously (*29*) on the backs of mice. Most mice reject epicutaneous tumors and the skin is populated with antitumor T_RM_ cells. For flow cytometry single cell suspensions were reconstituted in PBS and stained with live/dead discrimination dye (BioLegend) for 10 min at room temperature. Cell were washed with FACS buffer (PBS supplemented with 2% FBS, 2 mM EDTA, and 0.001% NaN_3_) and stained with H-2K^b^/SIINFEKL-pentamer (ProImmune) in FACS buffer for 10 min at room temperature, followed by staining with the remaining antibody mix for 20 min on ice. After staining, cell were washed with FACS buffer and fixed with 200 ul paraformaldehyde (BioLegend). All flow cytometric data acquisition was conducted on a Cytek Aurora within 24 hours after fixation.

### Imaging (H&E and tissue-based CyCIF)

H&E-stained FFPE sections were imaged using CyteFinder slide scanning fluorescence microscope (RareCyte Inc.) with a 20×/0.75 NA objective with no pixel binning. Serial FFPE sections (5 μm thick) were subjected to whole-slide CyCIF imaging with respective antibody panels for human and mouse tissue (Supplementary Table S1). CyCIF was performed as described in (PMID:29993362) and specifically as in protocols.io (dx.doi.org/10.17504/protocols.io.j8nlkoqbdv5r/v1). In brief, FFPE slides were baked using the BOND RX Automated IHC Stainer at 60°C for 30 minutes, dewax using Bond Dewax solution at 72°C, and antigen retrieval was performed using Epitope Retrieval 1 (Leica) solution at 100°C for 20 minutes. Multiple cycles of antibody incubation (overnight at 4°C in the dark), imaging, and fluorophore inactivation performed as cyclic imaging process. A 0.15mm single-sided self-adhesive spacer was applied to the slide followed by wet-mounting of a glass coverslip using 50% glycerol in 1× PBS for human tissue sections. Mouse tissue sections were imaged without spacers. Images were acquired using CyteFinder slide scanning fluorescence microscope (RareCyte Inc.) with a 20×/0.75 NA objective with no pixel binning. Coverslips were removed by soaking the slides in 42°C PBS and fluorophores were bleached by incubating slides in a solution of 4.5% H_2_O_2_ and 24 mmol/L NaOH in PBS and placing them under an LED light source for 1 hour. For mouse tissues were bleached twice for 45 mins in the same bleaching solution under LED light source. The list of all antibody panels used for both human and mouse tissue imaging are listed in Supplementary Table S1. Antibodies that passed a multi-step validation process and followed the expected staining pattern were included for downstream analysis.

### CyCIF image pre-processing and quality control

MCMICRO pipeline (RRID:SCR_022832), an open-source multiple-choice microscopy pipeline (version:38182748aa0ec021f684ce47248c57340d2f4cc7; full codes available on at https://github.com/labsyspharm/mcmicro) was used to stitch individual CyCIF images together into a high- dimensional representation for further segmentation and analyses. Specific parameters used were optimized after iterative inspection of results, specifically focused on performance of the segmentation module to ensure accurate identification of single cells (params.yml files available at https://github.com/labsyspharm/2025-Vallius-Shi-Novikov-melanoma-PCAII). The mean fluorescence intensities of each marker for each cell were computed after generating the segmentation masks resulting in a single-cell data table for each acquired whole-slide CyCIF image. Several steps were also taken to ensure the quality of the single-cell data. At the image level, the cross-cycle image registration and tissue integrity were reviewed. Regions with poor registration, tissues deformity, or other artifacts were identified and excluded from downstream single cell analysis. Antibodies with low confidence staining patterns were also excluded from the analyses. Segmentation parameters were iteratively changed to improve the accuracy of the segmentation masks.

### CyCIF single-cell phenotyping

As previously described (PMID:29993362), gating-based phenotyping approach was used to classify cells using an open-source visual gating tool (https://github.com/labsyspharm/gater) to determine gates for each marker. The identified gates for each marker were then used to rescale the single-cell data between 0 and 1, such that the values above 0.5 identify cells that express the marker. SCIMAP (PMID:38873023) python package (RRID:SCR_024751) were used to for cell-type calling based on a hierarchical classification. The assigned cell types were verified by overlaying the phenotypic labels onto the images.

### Spatial Analysis

Distance analyses were performed using the functions spatial_count and spatial_distance within the SCIMAP(PMID:38873023) python package. To determine the spatial relationship within the CYCIF data, average shortest distances between selected cell types were compute from the cell centroid as measured by the Euclidean distance between X/Y coordinates. The spatial distances were computed for selected cell types using spatial_distance. Subregions with a thickness of subsequent 800-micron starting from epidermis were identified using an open-source software package SpatialsCells (PMID:38701421).

### Statistical Tests

All statistical analysis was done in GraphPad Prism (GraphPad Software Inc.). In all experiments, p<0.05 was considered significant. Statistical details for experiments are provided in the figure legends. Box plots show 25th to 75th percentiles, with the median as the center and the whiskers corresponding to the minimum and maximum values. For CyCIF, statistical tests to infer P value for significant differences in mean were performed using the Mann-Whitney U rank test with mann whitney function in the scipy Python package.

## Acknowledgments

We want to thank the Center for Comparative Medicine at Massachusetts General Hospital for animal caretaking, the Flow Cytometry Core at Beth Israel Deaconess Medical Center for cell sorting assistance, and the Molecular Biology Core Facilities at Dana-Farber Cancer Institute for RNA sequencing. This work was made possible by the generous support from the American Cancer Society Postdoctoral Fellowship (PF-22-087-01-IBCD to J.B.W.), NIH training grant (T32AR007098 to J.B.W.), and the National Institute of Health (R01AR065807, R01CA279457, R01AI175582, R01AI127654 to TSK). P.K.S. is a cofounder andmember of the Board of Directors of Glencoe Software and a member of the Scientific Advisory Board for RareCyte and Montai Health; he holds equity in Glencoe and RareCyte and is also a consultant for Merck. This work was supported by the Harvard Ludwig Center, and an ASPIRE Award from The Mark Foundation for Cancer Research. SMP is supported by postdoctoral fellowship from Sigrid Juselius Foundation and Instrumentarium Foundation.

## Key Resources Table

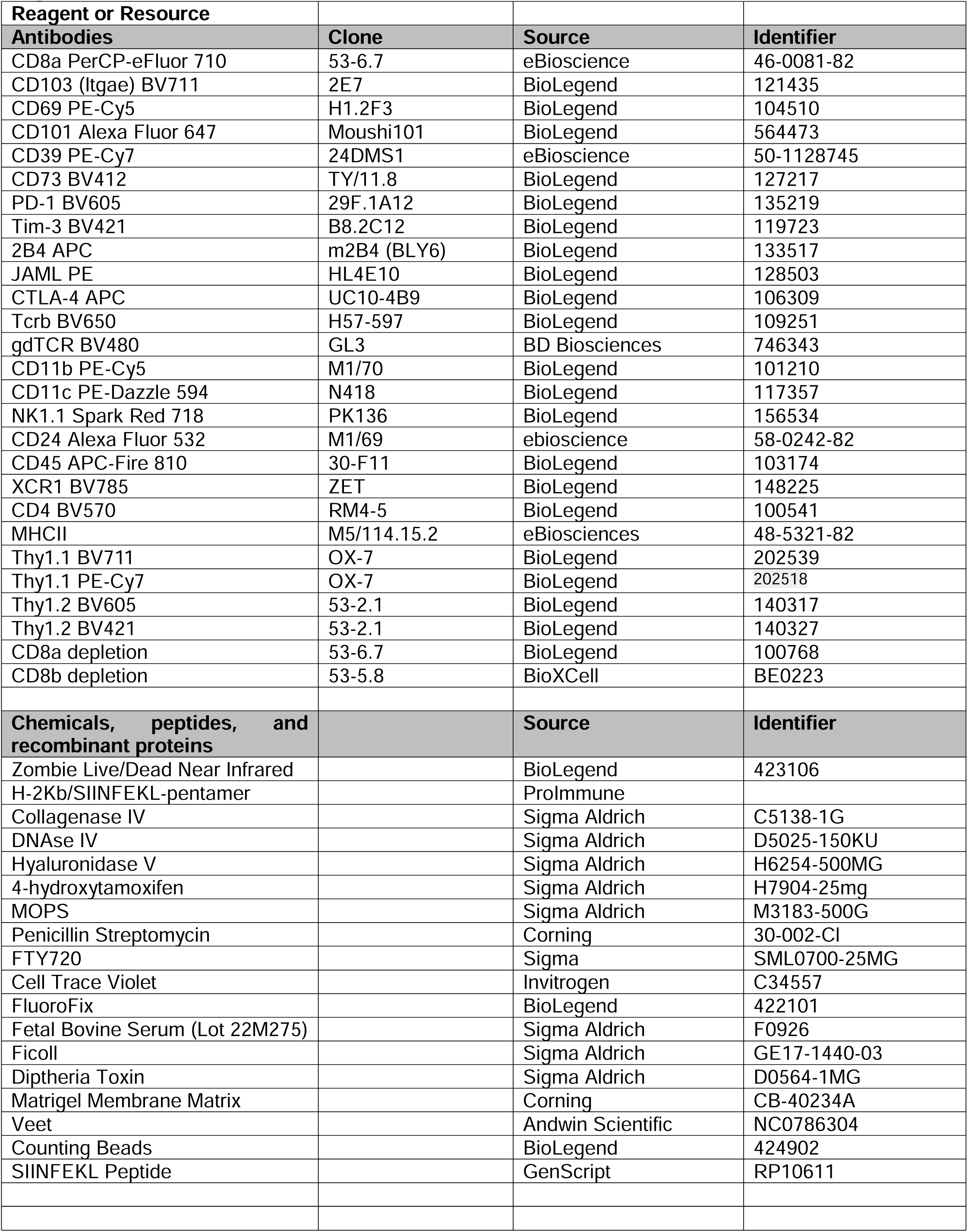
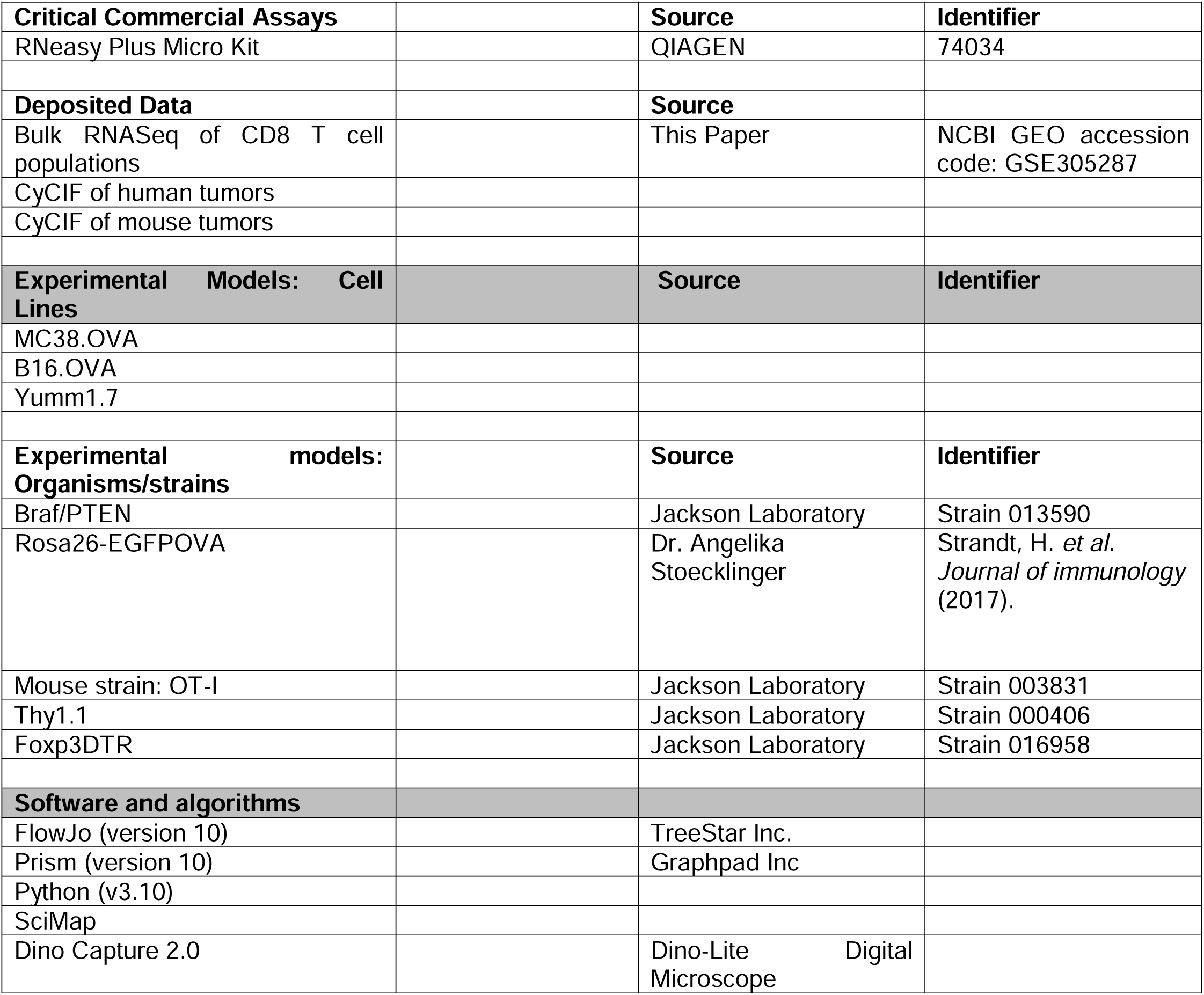

---

**Supplementary Figure S1.**
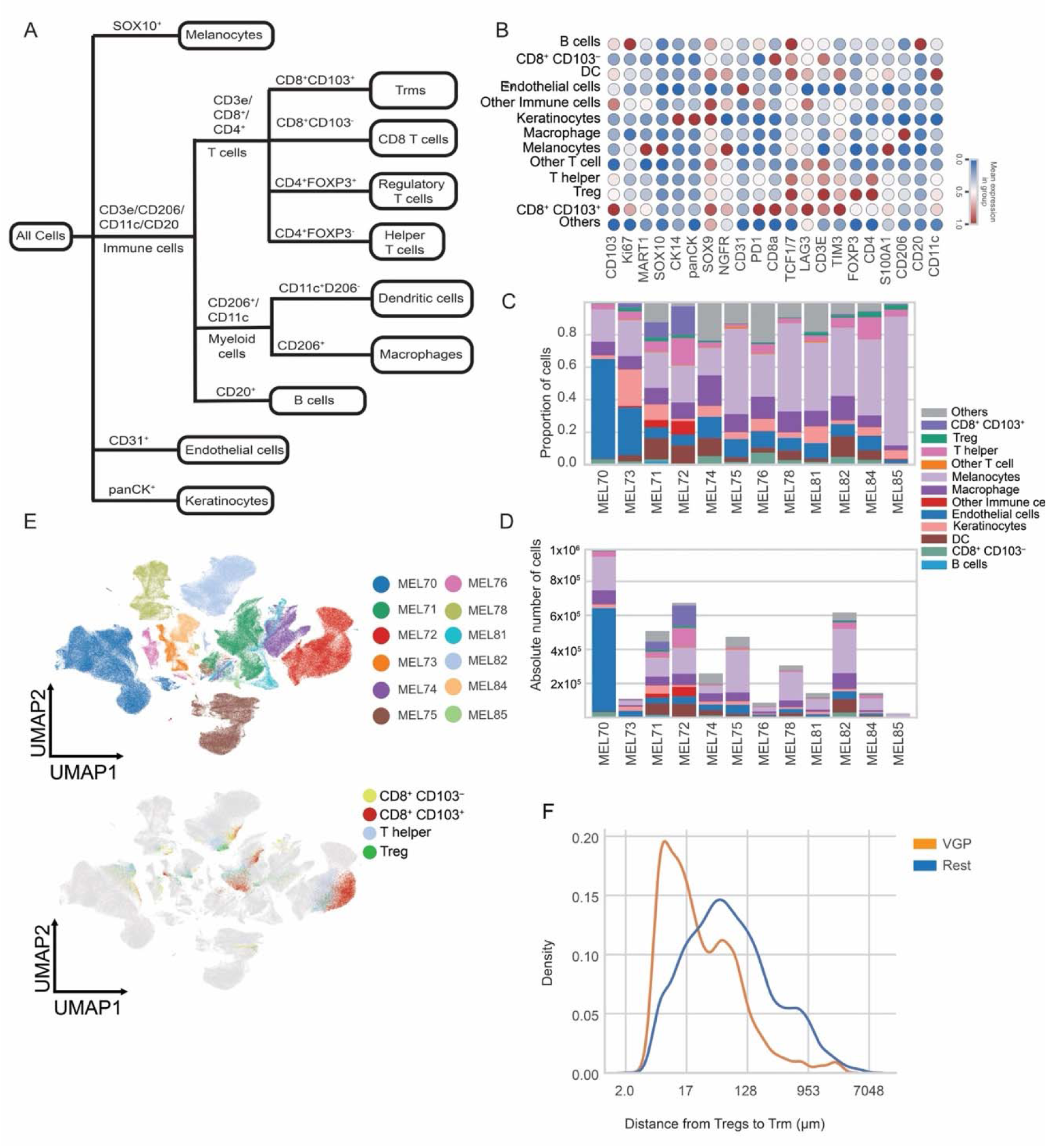
Multiplex imaging panel validation and immune cell composition in human stage II melanoma. **A**, Overview of the 21-marker t-CyCIF antibody panel used to identify immune and stromal cell populations, annotated by lineage. **B**, Heatmap showing scaled (z-score) expression of lineage and functional markers across single-cell clusters identified from multiplex imaging data. CD8^+^ T cells are stratified by CD103 expression, and CD4^+^ T cells by Foxp3 expression. **C** and **D**, Relative proportions (**C**) and absolute cell numbers (**D**) of immune, stromal, and melanoma cell populations across all samples. **E**, UMAP plots colored by samples and CD8^+^CD103^−^, CD8^+^CD103^+^, CD4^+^Foxp3^−^, and CD4^+^Foxp3^+^ T cell populations. **F**, Distribution plot showing Treg to CD8^+^CD103^+^ T cell distances. Significance was determined by a Mann-Whitney U rank test was used in **F**.

**Supplementary Figure S2.**
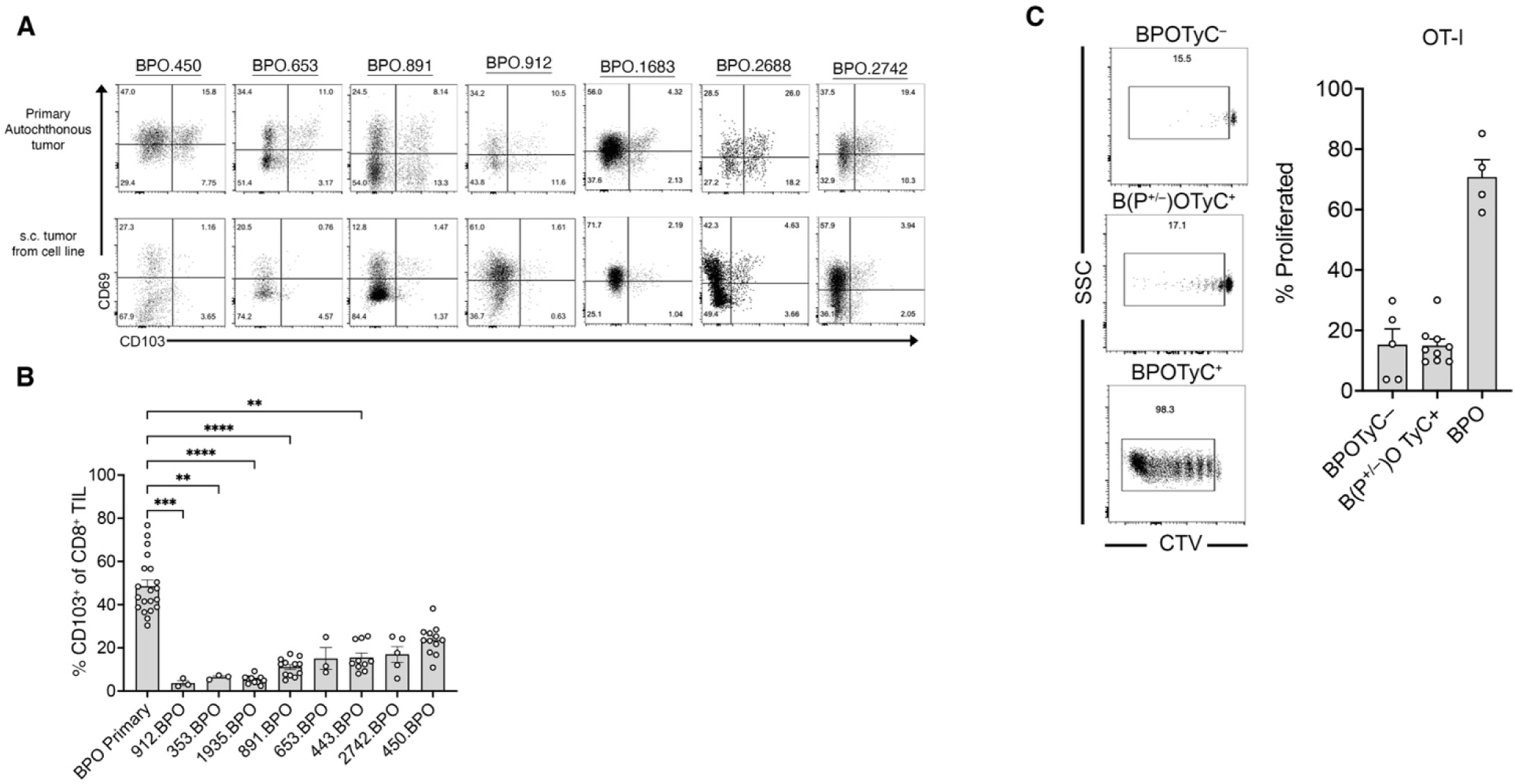
T_RM_-like CD8^+^ TIL frequencies in engraftable versus autochthonous melanoma models and OT-I cell proliferation. **A**, Representative flow cytometry plots of CD8 TILs expressing CD103 and CD69 in tumors from BPO-derived cell lines engrafted s.c. and their corresponding primary autochthonous BPO tumors. **B**, Average frequencies of T_RM_-like CD8^+^ TILs in primary autochthonous tumors versus s.c. engrafted tumors. BPO primary (autochthonous) tumors were taken at the time of s.c. engraftable experiments and are not the primary BPO-derived cell line tumors. **C**, OT-I proliferation in TdLN after transfer into tumor bearing or tumor-free BPO mice. Graphs display all mice from two (**A** and **B**) or one (**C**) independent experiment. Significance was determined by a Kruskal-Wallis test with Dunn’s multiple comparison correction in **B**.

**Supplementary Figure S3.**
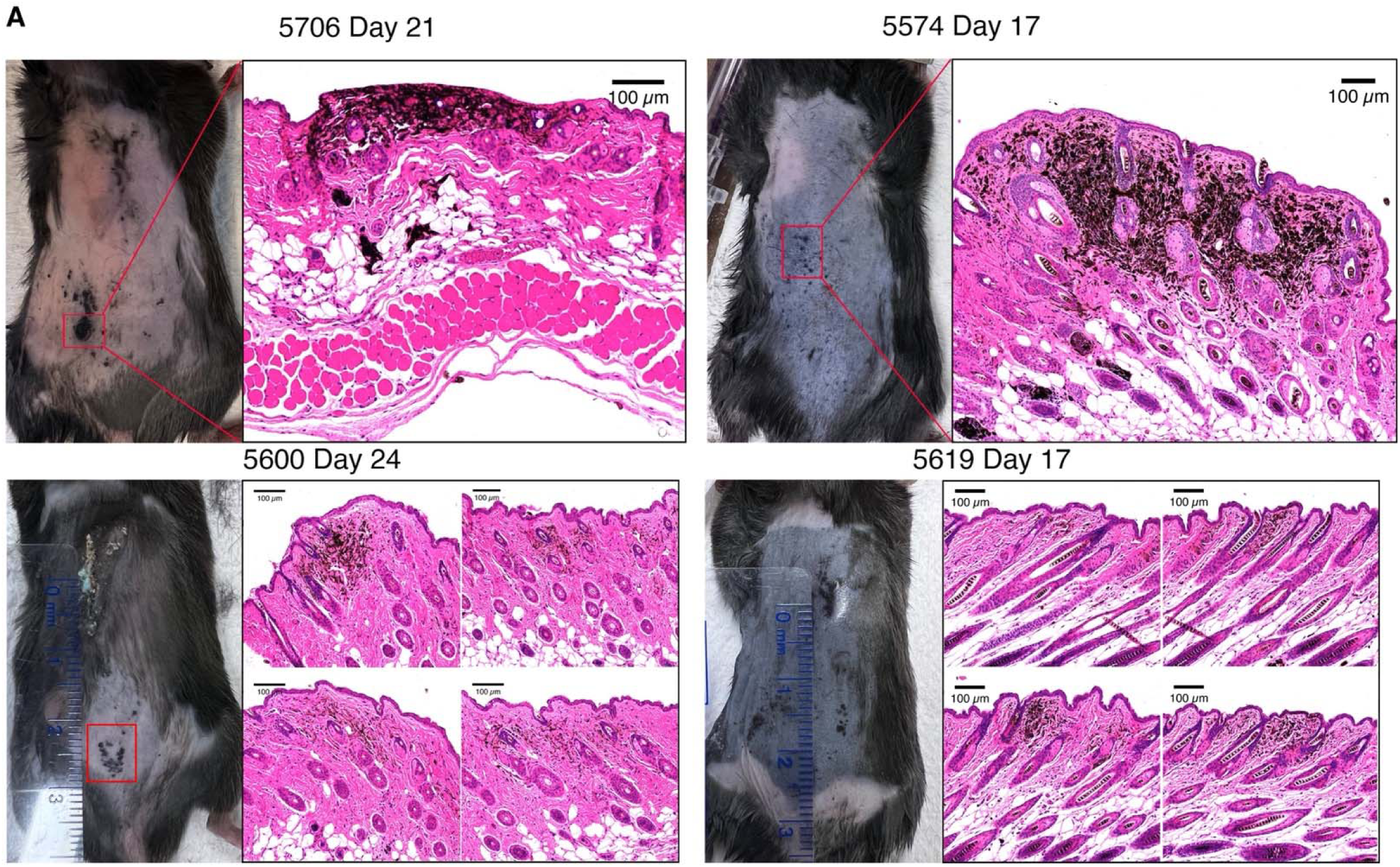
Histopathology of early-stage tumors. **A**, Representative H&E images showing hyperpigmented lesions confined to upper dermis.

**Supplementary Figure S4.**
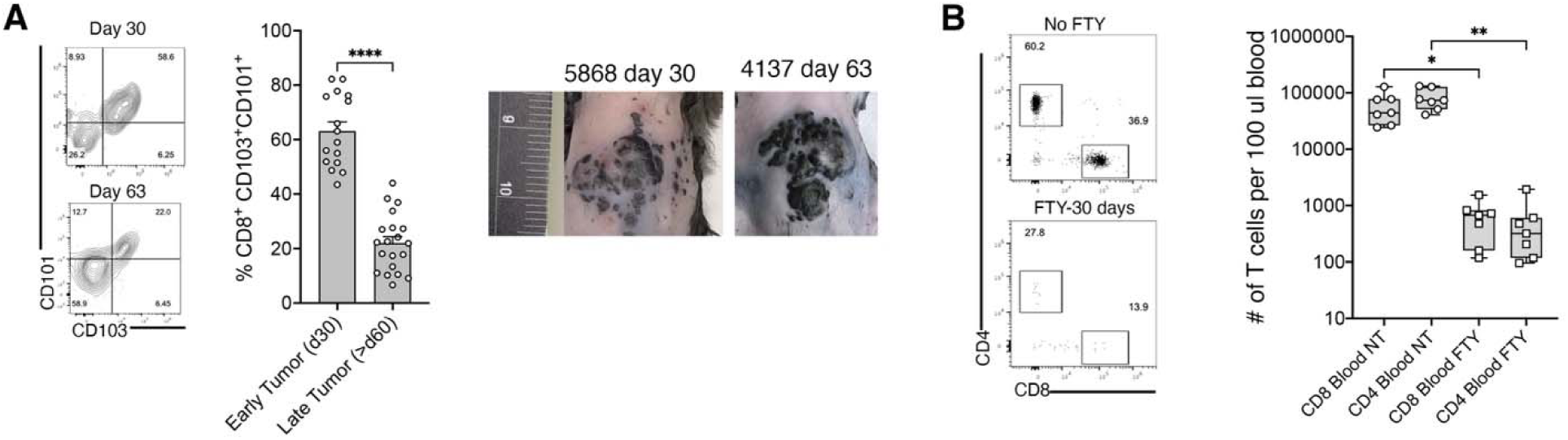
Stability of T_RM_-like CD8^+^ TIL niche during tumor progression. **A**, Representative flow cytometry plots showing CD103 and CD101 expression in early- and late-stage tumors. **B**, Representative flow cytometry plots and cell counts of T cells in peripheral blood after 30 days of FTY720 treatment. Graphs display all mice from 2 independent experiments. Significance was determined by a Mann-Whitney U rank test in **A** and a Kruskal-Wallis test with Dunn’s multiple comparison correction in **B.**

**Supplementary Figure S5.**
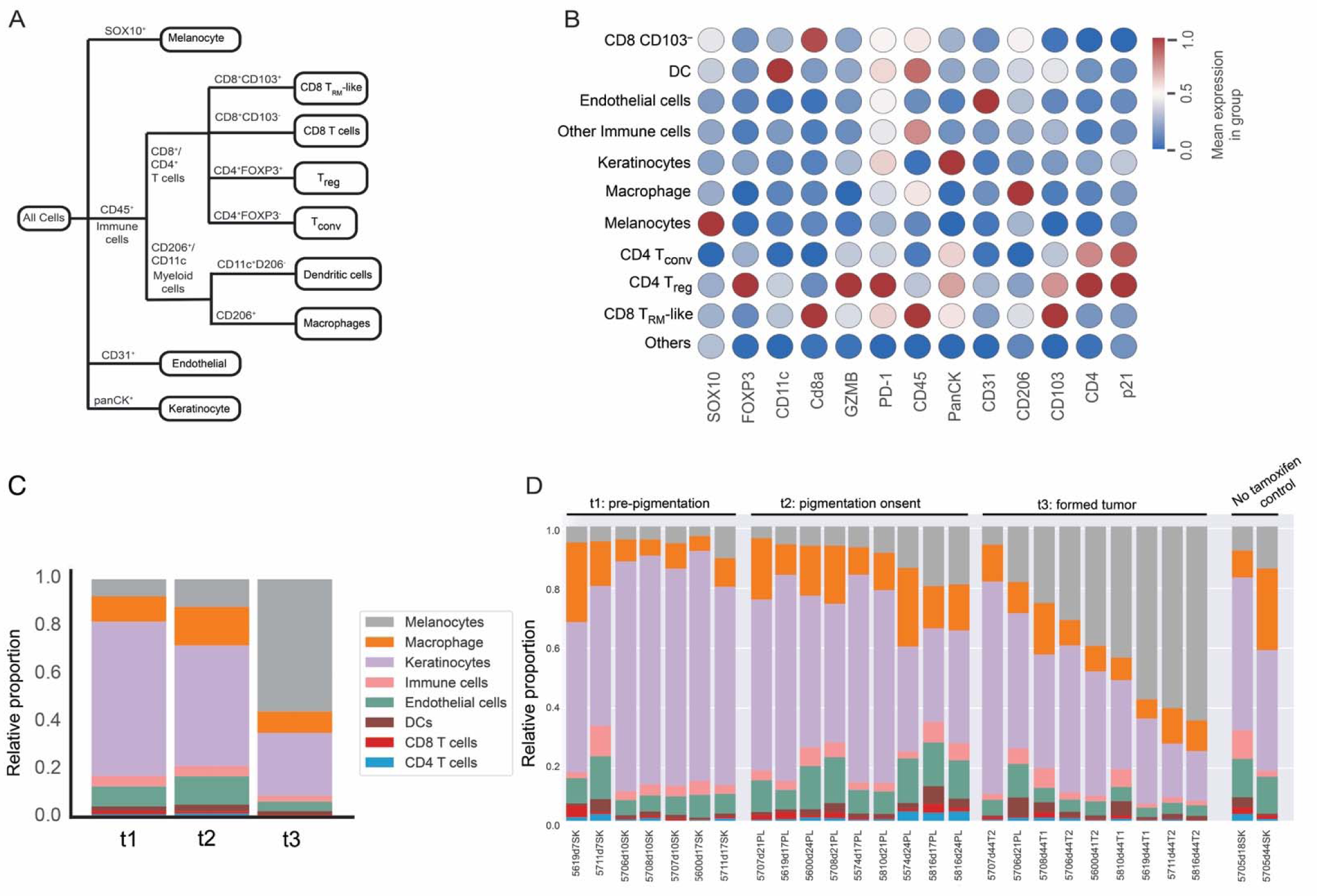
Immune-focused t-CyCIF antibody panel and melanoma biopsy quantification. **A**, Antibody panel composition for immune cell subset identification in mouse t-CyCIF. **B**, Expression of markers for cell phenotyping. **C** and **D,** Relative proportion of immune cell and stromal cell subsets across time points (**C**) or individual mice (**D**).

**Supplementary Figure S6.**
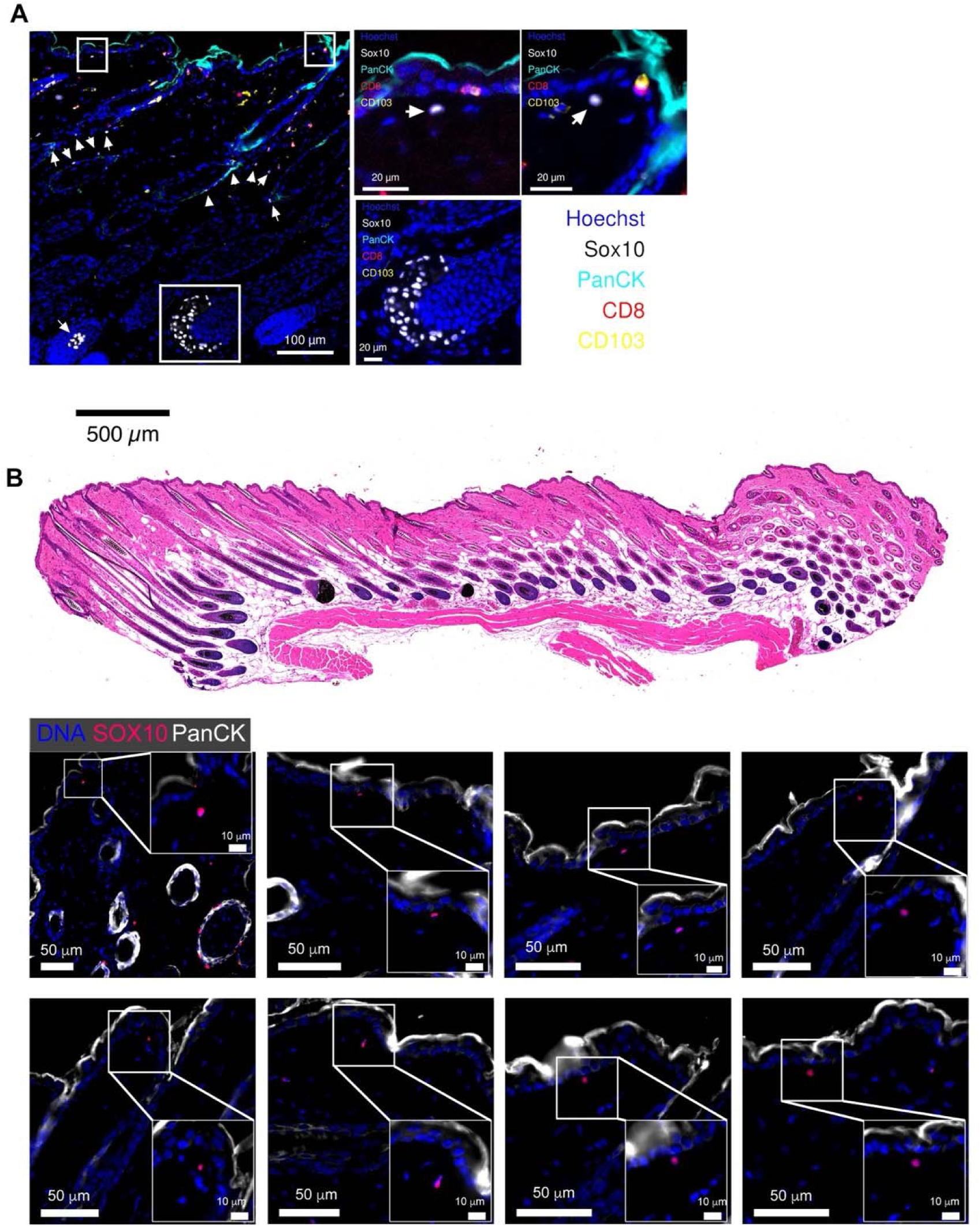
Sox10^+^ melanoma cell localization in pre-pigmentation stage. **A**, Representative t-CyCIF image showing Sox10^+^ melanoblast localization to lower hair follicle bulb. **B**, Scattered dermal and papillary dermis Sox10^+^ melanoma cells (white arrows) in pre-pigmentation biopsies.

**Supplementary Figure S7.**
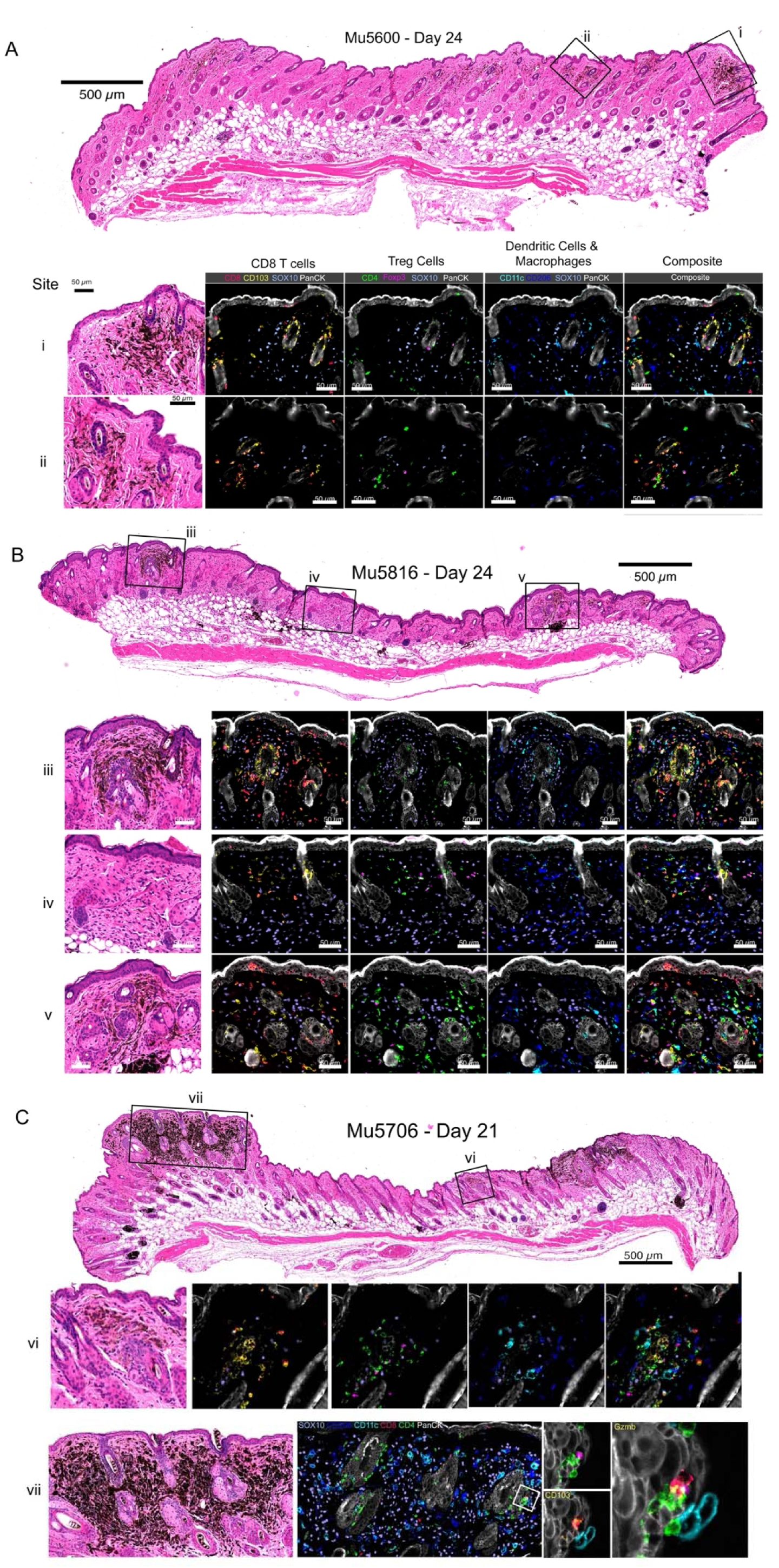
Heterogeneity of immune microenvironments in early-stage hyperpigmented melanoma cell clusters. **A**-**C**, Representative H&E and t-CyCIF images highlighting immune cell composition in hyperpigmented lesions. Sites i-iv show the presence of T_RM_-like cells clustered with Sox10^+^ melanoma cells. Sites iv-vi exemplify Treg cell enrichment in some hyperpigmented areas. Sites iv and vii, display macrophage-dominant clusters with sparse CD8^+^ T cells. Inset in site vii shows a single Gzmb^+^ T_RM_-like cell in contact with a Treg cell.

**Supplementary Figure S8.**
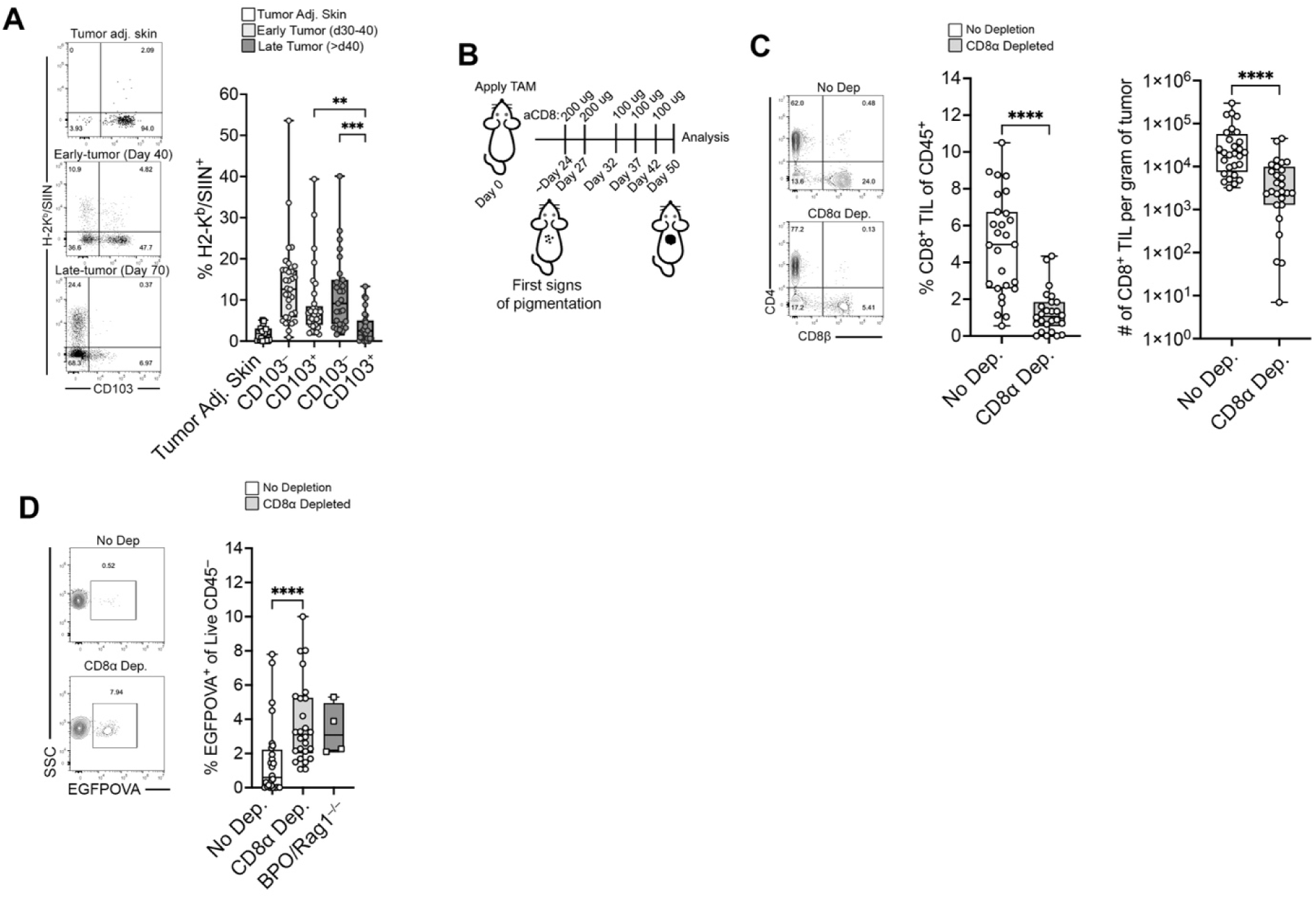
CD8^+^ T_RM_-like cell tumor specificity and role in immunoediting. **A**, H-2K^b^/SIINFEKL pentamer staining showing tumor antigen specificity in T_RM_-like versus CD103^−^ CD8^+^ TILs at early and late stages of tumor development. **B**, Experimental schematic for CD8^+^ T cell depletion starting at pigmentation onset. **C**, Confirmation of CD8^+^ TIL depletion at endpoint. **D**, Frequency of EGFPOVA^+^ tumor cells in nontreated and CD8-depleted versus immunodeficient (BPO/Rag1^−/–^) mice. Significance was determined by a Kruskal-Wallis test with Dunn’s multiple comparison correction in **A** and **D** and a Mann-Whitney U rank test in **C**.

**Supplementary Figure S9.**
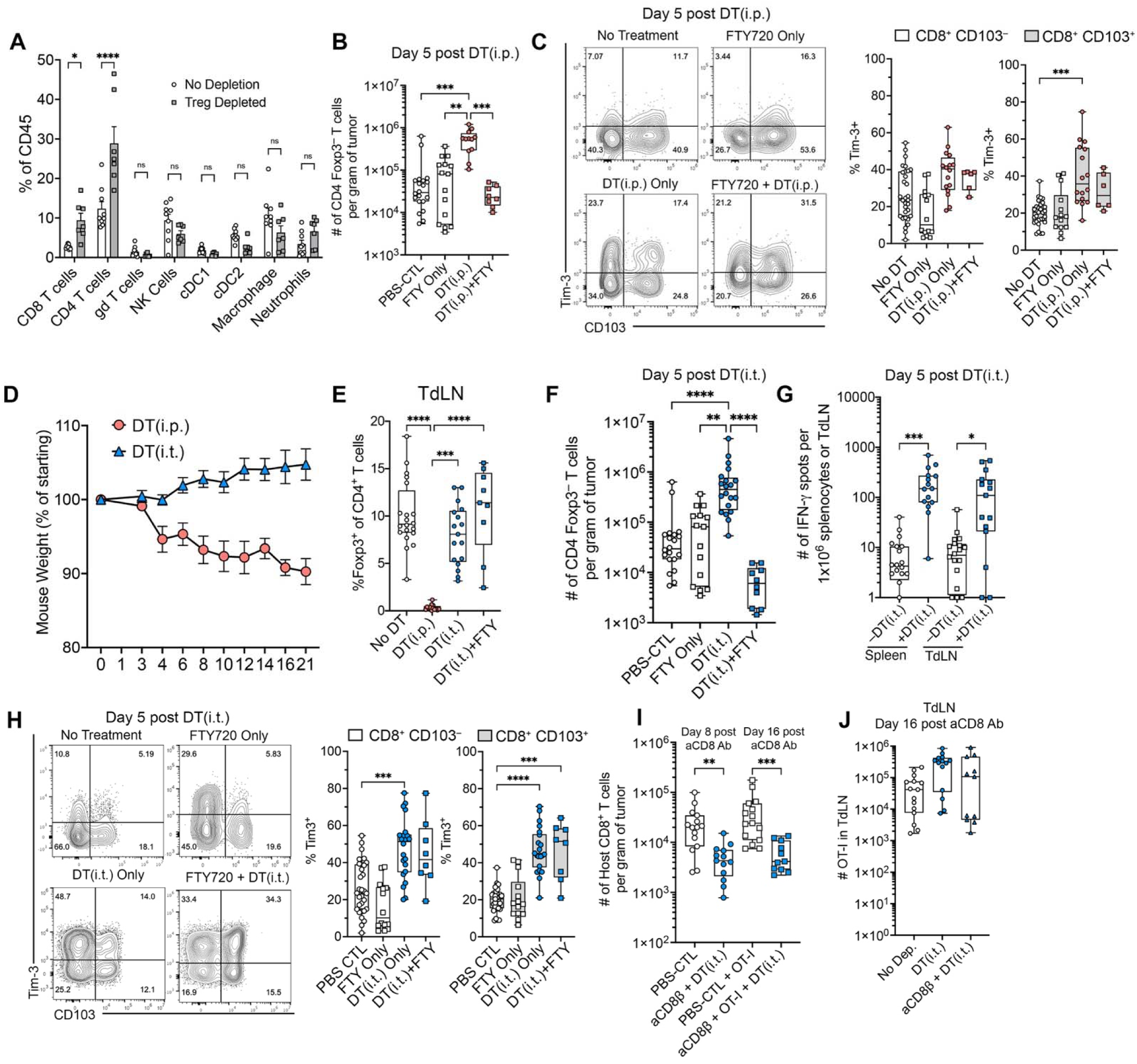
Tumor immune compartment changes following systemic and local Treg depletion, and confirmation of CD8^+^ T cell depletion. **A**, Frequency of CD8^+^ T cells, CD4^+^ T cells, γδ T cells, NK cells, cDC1, cDC2, macrophages, and neutrophils among CD45^+^ cells in the tumor at day 5 following systemic Treg depletion (DT i.p.) versus no depletion controls. **B**, Number of CD4^+^ Foxp3^−^ Tconv cells per gram of tumor on day 5 following systemic Treg depletion (DT i.p.) ± FTY720. **C**, Representative flow cytometry plots and quantification of Tim-3 expression on CD8^+^ CD103^−^ and CD8^+^ CD103^+^ TIL subsets on day 5 following DT(i.p.) ± FTY720. **D**, Mouse weight (% of starting weight) over time following systemic (DT i.p. on day 0 and 1) versus intratumoral (DT i.t. once every 7 days) Treg depletion. **E**, Frequency of Foxp3^+^ cells among CD4^+^ T cells in the TdLN following no depletion, DT(i.p.), DT(i.t.), or DT(i.t.) + FTY720. **F**, Number of CD4^+^ Foxp3^−^ Tconv cells per gram of tumor on day 5 following intratumoral Treg depletion (DT i.t.) ± FTY720. **G**, Number of IFN-γ spots per 1 × 10^6^ splenocytes or TdLN cells in response to SIINFEKL peptide stimulation on day 5 following DT(i.t.) versus no DT controls. **H**, Representative flow cytometry plots and quantification of Tim-3 expression on CD8^+^ CD103^−^ and CD8^+^ CD103^+^ TIL subsets on day 5 following DT(i.t.) ± FTY720. **I**, Number of host CD8^+^ T cells per gram of tumor at day 8 (Tconv recruitment experiment, corresponding to Figure 7G) and day 16 (OT-I recruitment experiment, corresponding to Figure 7H) after anti-CD8β antibody administration, confirming sustained CD8^+^ T cell reduction at the time of analysis. **J**, Number of OT-I T cells in the TdLN at day 16 post anti-CD8β antibody administration, in mice receiving no depletion (PBS-treated), DT(i.t.), or anti-CD8β + DT(i.t.). Graphs display all mice from 2–5 independent experiments. Significance was determined by a Mann-Whitney U test in **A** and **G**, and by a Kruskal-Wallis test with Dunn’s multiple comparison correction in **B**, **C**, **E**, **F**, **H**, **I**, and **J**.

**Supplementary Figure S10.**
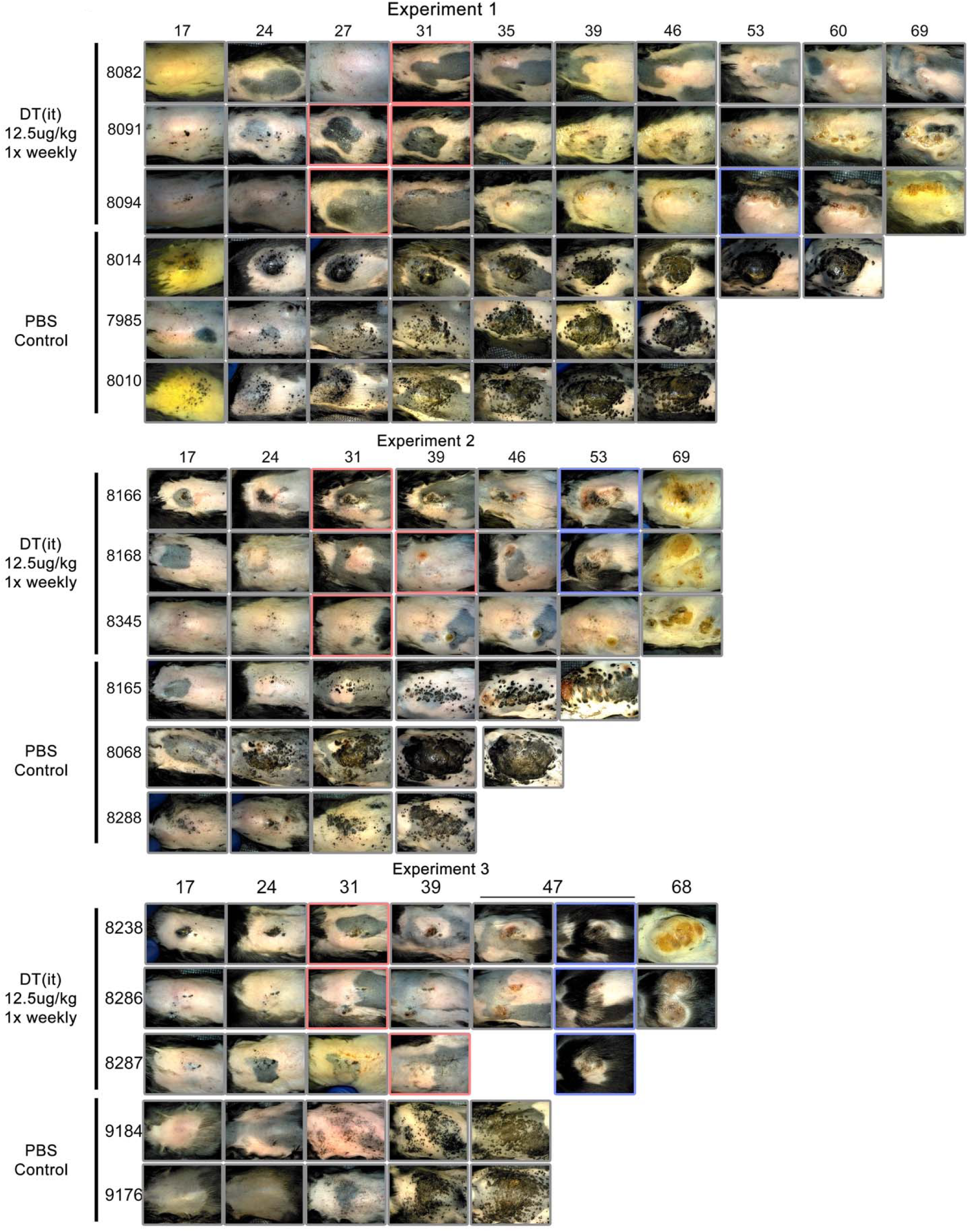
Serial digital microscopy of tumor lesions following weekly intratumoral Treg depletion. Representative images captured by handheld digital microscopy (DinoLite) of BPO tumor lesions over time in mice receiving weekly intratumoral DT (DT(i.t.), 12.5 µg/kg, once per week) or PBS-treated controls, shown across three independent experiments. Each row represents an individual mouse (mouse ID indicated at left), and each column corresponds to a single timepoint (days after tamoxifen application, indicated at top). Red boxes highlight timepoints at which visible diminishment of pigmented lesions was observed in DT(i.t.)-treated mice. Blue boxes highlight timepoints at which vitiligo became apparent at the treatment site. PBS-treated controls did not exhibit lesion regression or vitiligo at any timepoint examined. Animals were removed from the study upon reaching endpoint criteria, accounting for incomplete time series in some rows.

**Supplementary Figure S11.**
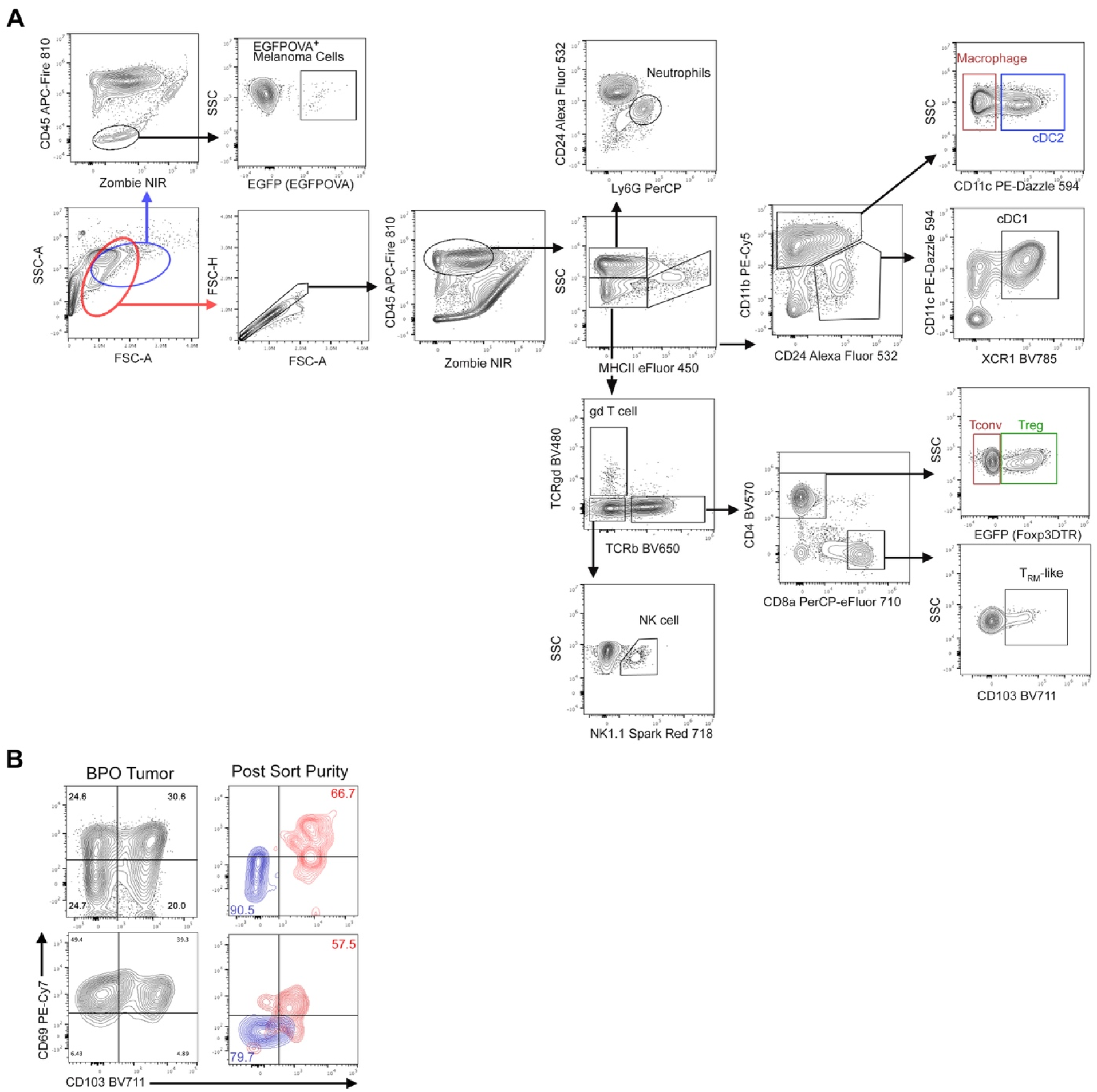
Flow cytometry gating strategy. **A**, Separate gates were used to isolate lymphocytes and tumor cells. Tumor cells were identified to be within the CD45^−^ SSChi fraction. Cell populations were identified as follows; Neutrophils: CD45^+^ SSChi MHCII^−^ CD24int Ly6G^+^, Macrophage: CD45^+^ MHCII^+^ CD11b^+^ CD24^−^ CD11c^−^, cDC1: CD45^+^ MHCII^+^ CD24^+^ CD11b^−^ CD11c^+^ XCR1^+^, gd T cell: CD45+ SSClow TCRb^−^ TCRgd^+^, NK cell: CD45^+^ SSClow TCRb^−^ TCRgd^−^ NK1.1^+^, CD8 T cell: CD45^+^ SSClow TCRb^+^ CD8a^+^, CD4 T cell: CD45^+^ SSClow TCRb^+^ CD4^+^. Treg cells were identified as CD4^+^ T cell with Foxp3EGFPDTR^+^. TRM-like cells were identified as CD8^+^ T cell with CD103 expression. **B**, Mid-sort analysis for bulk RNASeq experiment. Cell purity for two samples was determined during the sort to confirm purity.

## REFERENCES

1. P. Sharma, S. Hu-Lieskovan, J. A. Wargo, A. Ribas, Primary, Adaptive, and Acquired Resistance to Cancer Immunotherapy. Cell 168, 707–723 (2017).

2. A. Ribas, J. D. Wolchok, Cancer immunotherapy using checkpoint blockade. Science 359, 1350–1355 (2018).

3. G. P. Dunn, L. J. Old, R. D. Schreiber, The Three Es of Cancer Immunoediting. Annual Review of Immunology 22, 329–360 (2004).

4. C. M. Koebel, W. Vermi, J. B. Swann, N. Zerafa, S. J. Rodig, L. J. Old, M. J. Smyth, R. D. Schreiber, Adaptive immunity maintains occult cancer in an equilibrium state. Nature 450, 903–907 (2007).

5. S. N. Mueller, L. K. Mackay, Tissue-resident memory T cells: local specialists in immune defence. Nature Reviews Immunology 16, 79–89 (2016).

6. N. V. Gavil, K. Cheng, D. Masopust, Resident memory T cells and cancer. Immunity 57, 1734–1751 (2024).

7. Y. Pan, T. Tian, C. O. Park, S. Y. Lofftus, S. Mei, X. Liu, C. Luo, J. T. O’Malley, A. Gehad, J. E. Teague, S. J. Divito, R. Fuhlbrigge, P. Puigserver, J. G. Krueger, G. S. Hotamisligil, R. A. Clark, T. S. Kupper, Survival of tissue-resident memory T cells requires exogenous lipid uptake and metabolism. Nature 543, 252–256 (2017).

8. N. P. Smith, Y. Yan, Y. Pan, J. B. Williams, K. Manakongtreecheep, S. M. Pant, J. Zhao, T. Tian, T. Pan, b C. Stingley, K. Wu, J. Zhang, A. L. Kley, P. K. Sorger, A.-C. Villani, T. S. Kupper, Resident memory T cell development is gradual and shows AP-1 gene expression in mature cells. JCI Insight 10, e187381 (2025).

9. M. Evrard, E. Becht, R. Fonseca, A. Obers, S. L. Park, N. Ghabdan-Zanluqui, J. Schroeder, S. N. Christo, D. Schienstock, J. Lai, T. N. Burn, A. Clatch, I. G. House, P. Beavis, A. Kallies, F. Ginhoux, S. N. Mueller, R. Gottardo, E. W. Newell, L. K. Mackay, Single-cell protein expression profiling resolves circulating and resident memory T cell diversity across tissues and infection contexts. Immunity, doi: 10.1016/j.immuni.2023.06.005 (2023).

10. M. C. Scott, Z. Steier, M. J. Pierson, J. M. Stolley, S. D. O’Flanagan, A. G. Soerens, S. P. Wijeyesinghe, L. K. Beura, G. Dileepan, B. J. Burbach, M. Künzli, C. F. Quarnstrom, O. C. G. Smith, E. Weyu, S. E. Hamilton, V. Vezys, A. K. Shalek, D. Masopust, Deep profiling deconstructs features associated with memory CD8+ T cell tissue residence. Immunity 58, 162–181.e10 (2025).

11. S. Ariotti, M. A. Hogenbirk, F. E. Dijkgraaf, L. L. Visser, M. E. Hoekstra, J.-Y. Song, H. Jacobs, J. B. Haanen, T. N. Schumacher, Skin-resident memory CD8^+^ T cells trigger a state of tissue-wide pathogen alert. Science 346, 101–105 (2014).

12. J. M. Schenkel, K. A. Fraser, V. Vezys, D. Masopust, Sensing and alarm function of resident memory CD8^+^ T cells. Nature Immunology 14, 509–513 (2013).

13. E. Menares, F. Gálvez-Cancino, P. Cáceres-Morgado, E. Ghorani, E. López, X. Díaz, J. Saavedra-Almarza, D. A. Figueroa, E. Roa, S. A. Quezada, A. Lladser, Tissue-resident memory CD8+ T cells amplify anti-tumor immunity by triggering antigen spreading through dendritic cells. Nature Communications 10, 4401 (2019).

14. J. M. Schenkel, K. A. Fraser, L. K. Beura, K. E. Pauken, V. Vezys, D. Masopust, Resident memory CD8 T cells trigger protective innate and adaptive immune responses. Science 346, 1254536–101 (2014).

15. L. R. Shiow, D. B. Rosen, N. Brdičková, Y. Xu, J. An, L. L. Lanier, J. G. Cyster, M. Matloubian, CD69 acts downstream of interferon-α/β to inhibit S1P1 and lymphocyte egress from lymphoid organs. Nature 440, 540–544 (2006).

16. L. K. Mackay, A. Rahimpour, J. Z. Ma, N. Collins, A. T. Stock, M.-L. Hafon, J. Vega-Ramos, P. Lauzurica, S. N. Mueller, T. Stefanovic, D. C. Tscharke, W. R. Heath, M. Inouye, F. R. Carbone, T. Gebhardt, The developmental pathway for CD103(+)CD8+ tissue-resident memory T cells of skin. Nature Immunology 14, 1294–1301 (2013).

17. R. Watanabe, A. Gehad, C. Yang, L. L. Scott, J. E. Teague, C. Schlapbach, C. P. Elco, V. Huang, T. R. Matos, T. S. Kupper, R. A. Clark, Human skin is protected by four functionally and phenotypically discrete populations of resident and recirculating memory T cells. Science Translational Medicine 7, 279ra39–279ra39 (2015).

18. J. Edwards, J. S. Wilmott, J. Madore, T. N. Gide, C. Quek, A. Tasker, A. Ferguson, J. Chen, R. Hewavisenti, P. Hersey, T. Gebhardt, W. Weninger, W. J. Britton, R. P. M. Saw, J. F. Thompson, A. M. Menzies, G. V. Long, R. A. Scolyer, U. Palendira, CD103+ Tumor-Resident CD8+ T Cells Are Associated with Improved Survival in Immunotherapy-Naïve Melanoma Patients and Expand Significantly During Anti–PD-1 Treatment. Clinical cancer research : an official journal of the American Association for Cancer Research 24, 3036–3045 (2018).

19. A.-P. Ganesan, J. Clarke, O. Wood, E. M. Garrido-Martin, S. J. Chee, T. Mellows, D. Samaniego-Castruita, D. Singh, G. Seumois, A. Alzetani, E. Woo, P. S. Friedmann, E. V. King, G. J. Thomas, T. Sanchez-Elsner, P. Vijayanand, C. H. Ottensmeier, Tissue-resident memory features are linked to the magnitude of cytotoxic T cell responses in human lung cancer. Nature Publishing Group 122, 3491 (2017).

20. J. Clarke, B. Panwar, A. Madrigal, D. Singh, R. Gujar, O. Wood, S. J. Chee, S. Eschweiler, E. V. King, A.S. Awad, C. J. Hanley, K. J. McCann, S. Bhattacharyya, E. Woo, A. Alzetani, G. Seumois, G. J. Thomas, A.-P. Ganesan, P. S. Friedmann, T. Sanchez-Elsner, F. Ay, C. H. Ottensmeier, P. Vijayanand, Single-cell transcriptomic analysis of tissue-resident memory T cells in human lung cancer. The Journal of experimental medicine 216, 2128–2149 (2019).

21. J. R. Webb, K. Milne, P. Watson, R. J. Deleeuw, B. H. Nelson, Tumor-infiltrating lymphocytes expressing the tissue resident memory marker CD103 are associated with increased survival in high-grade serous ovarian cancer. Clinical cancer research : an official journal of the American Association for Cancer Research 20, 434–444 (2014).

22. F. L. Komdeur, T. M. Prins, S. van de Wall, A. Plat, G. B. A. Wisman, H. Hollema, T. Daemen, D. N. Church, M. de Bruyn, H. W. Nijman, CD103+ tumor-infiltrating lymphocytes are tumor-reactive intraepithelial CD8+ T cells associated with prognostic benefit and therapy response in cervical cancer. Oncoimmunology 6, e1338230 (2017).

23. F. Djenidi, J. Adam, A. Goubar, A. Durgeau, G. Meurice, V. de Montpreville, P. Validire, B. Besse, F. Mami-Chouaib, CD8+CD103+ Tumor–Infiltrating Lymphocytes Are Tumor-Specific Tissue-Resident Memory T Cells and a Prognostic Factor for Survival in Lung Cancer Patients. Journal of immunology (Baltimore, Md. : 1950) 194, 3475–3486 (2015).

24. T. Murray, S. A. F. Marraco, P. Baumgaertner, N. Bordry, L. Cagnon, A. Donda, P. Romero, G. Verdeil, D. E. Speiser, Very Late Antigen-1 Marks Functional Tumor-Resident CD8 T Cells and Correlates with Survival of Melanoma Patients. Frontiers in immunology 7, 573 (2016).

25. I. Damei, T. Trickovic, F. Mami-Chouaib, S. Corgnac, Tumor-resident memory T cells as a biomarker of the response to cancer immunotherapy (2023). file:///Users/jbw38/Downloads/fimmu-14-1205984.pdf.

26. J. Han, Y. Zhao, K. Shirai, A. Molodtsov, F. W. Kolling, J. L. Fisher, P. Zhang, S. Yan, T. G. Searles, J. M. Bader, J. Gui, C. Cheng, M. S. Ernstoff, M. J. Turk, C. V. Angeles, Resident and circulating memory T cells persist for years in melanoma patients with durable responses to immunotherapy. Nat Cancer 2, 300–311 (2021).

27. C. Jiang, C.-C. Chao, J. Li, X. Ge, A. Shen, V. Jucaud, C. Cheng, X. Shen, Tissue-resident memory T cell signatures from single-cell analysis associated with better melanoma prognosis. iScience 27, 109277 (2024).

28. B. T. Malik, K. T. Byrne, J. L. Vella, P. Zhang, T. B. Shabaneh, S. M. Steinberg, A. K. Molodtsov, J. S. Bowers, C. V. Angeles, C. M. Paulos, Y. H. Huang, M. J. Turk, Resident memory T cells in skin mediate durable immunity to melanoma. Science immunology 2, eaam6346 (2017).

29. S. L. Park, A. Buzzai, J. Rautela, J. L. Hor, K. Hochheiser, M. Effern, N. McBain, T. Wagner, J. Edwards, R. McConville, J. S. Wilmott, R. A. Scolyer, T. T. ting, U. Palendira, D. Gyorki, S. N. Mueller, N. D. Huntington, S. Bedoui, M. H. lzel, L. K. Mackay, J. Waithman, T. Gebhardt, Tissue-resident memory CD8+ T cells promote melanoma–immune equilibrium in skin. Nature 565, 1–23 (2019).

30. C. Yapp, A. J. Nirmal, F. Zhou, A. Y. H. Wong, J. B. Tefft, Y. D. Lu, Z. Shang, Z. Maliga, P. M. Llopis, G. F. Murphy, C. G. Lian, G. Danuser, S. Santagata, P. K. Sorger, Highly Multiplexed 3D Profiling of Cell States and Immune Niches in Human Tumours. bioRxiv, 2023.11.10.566670 (2025).

31. H. Strandt, D. F. Pinheiro, D. H. Kaplan, D. Wirth, I. K. Gratz, P. Hammerl, J. Thalhamer, A. Stoecklinger, Neoantigen Expression in Steady-State Langerhans Cells Induces CTL Tolerance. Journal of immunology (Baltimore, Md. : 1950) 199, 1626–1634 (2017).

32. N. P. Smith, Y. Yan, Y. Pan, J. B. Williams, K. Manakongtreecheep, S. Pant, J. Zhao, T. Tian, T. Pan, C. Stingley, K. Wu, J. Zhang, A. L. Kley, P. K. Sorger, A.-C. Villani, T. S. Kupper, Resident memory T cell development is associated with AP-1 transcription factor upregulation across anatomical niches. bioRxiv, 2023.09.29.560006 (2023).

33. B. V. Park, Z. T. Freeman, A. Ghasemzadeh, M. A. Chattergoon, A. Rutebemberwa, J. Steigner, M. E. Winter, T. V. Huynh, S. M. Sebald, S.-J. Lee, F. Pan, D. M. Pardoll, A. L. Cox, TGFβ1-Mediated SMAD3 Enhances PD-1 Expression on Antigen-Specific T Cells in Cancer. Cancer Discov. 6, 1366–1381 (2016).

34. S. Terawaki, S. Chikuma, S. Shibayama, T. Hayashi, T. Yoshida, T. Okazaki, T. Honjo, IFN-α Directly Promotes Programmed Cell Death-1 Transcription and Limits the Duration of T Cell-Mediated Immunity. J. Immunol. 186, 2772–2779 (2011).

35. S. Spranger, D. Dai, B. Horton, T. F. Gajewski, Tumor-Residing Batf3 Dendritic Cells Are Required for Effector T Cell Trafficking and Adoptive T Cell Therapy. Cancer cell 31, 711–723.e4 (2017).

36. P. Savas, B. Virassamy, C. Ye, A. Salim, C. P. Mintoff, F. Caramia, R. Salgado, D. J. Byrne, Z. L. Teo, S. Dushyanthen, A. Byrne, L. Wein, S. J. Luen, C. Poliness, S. S. Nightingale, A. S. Skandarajah, D. E. Gyorki, C. M. Thornton, P. A. Beavis, S. B. Fox, K. C. F. C. for R. into F. B. C. (kConFab), P. K. Darcy, T. P. Speed, L. K. Mackay, P. J. Neeson, S. Loi, Single-cell profiling of breast cancer T cells reveals a tissue-resident memory subset associated with improved prognosis. Nature Medicine 24, 986–993 (2018).

37. S. G. D. León-Rodríguez, C. Aguilar-Flores, J. A. Gajón, Á. Juárez-Flores, A. Mantilla, R. Gerson-Cwilich, J. F. Martínez-Herrera, D. A. Villegas-Osorno, C. T. Gutiérrez-Quiroz, S. Buenaventura-Cisneros, M. A. Sánchez-Prieto, E. Castelán-Maldonado, S. R. Rivera, E. M. Fuentes-Pananá, L. C. Bonifaz, TCF1-positive and TCF1-negative TRM CD8 T cell subsets and cDC1s orchestrate melanoma protection and immunotherapy response. J. Immunother. Cancer 12, e008739 (2024).

38. A. M. Luoma, S. Suo, Y. Wang, L. Gunasti, C. B. M. Porter, N. Nabilsi, J. Tadros, A. P. Ferretti, S. Liao, C. Gurer, Y.-H. Chen, S. Criscitiello, C. A. Ricker, D. Dionne, O. Rozenblatt-Rosen, R. Uppaluri, R. I. Haddad, O. Ashenberg, A. Regev, E. M. V. Allen, G. MacBeath, J. D. Schoenfeld, K. W. Wucherpfennig, Tissue-resident memory and circulating T cells are early responders to pre-surgical cancer immunotherapy. Cell 185, 2918–2935.e29 (2022).

39. Y. Simoni, E. Becht, M. Fehlings, C. Y. Loh, S.-L. Koo, K. W. W. Teng, J. P. S. Yeong, R. Nahar, T. Zhang, H. Kared, K. Duan, N. Ang, M. Poidinger, Y. Y. Lee, A. Larbi, A. J. Khng, E. Tan, C. Fu, R. Mathew, M. Teo, W. T. Lim, C. K. Toh, B.-H. Ong, T. Koh, A. M. Hillmer, A. Takano, T. K. H. Lim, E.-H. Tan, W. Zhai, D. S. W. Tan, I. B. Tan, E. W. Newell, Bystander CD8+ T cells are abundant and phenotypically distinct in human tumour infiltrates. Nature 557, 575–579 (2018).

40. P. C. Rosato, S. Wijeyesinghe, J. M. Stolley, C. E. Nelson, R. L. Davis, L. S. Manlove, C. A. Pennell, B. R. Blazar, C. C. Chen, M. A. Geller, V. Vezys, D. Masopust, Virus-specific memory T cells populate tumors and can be repurposed for tumor immunotherapy. Nature Communications 10, 567 (2019).

41. W. Scheper, S. Kelderman, L. F. Fanchi, C. Linnemann, G. Bendle, M. A. J. de Rooij, C. Hirt, R. Mezzadra, M. Slagter, K. Dijkstra, R. J. C. Kluin, P. Snaebjornsson, K. Milne, B. H. Nelson, H. Zijlmans, G. Kenter, E. E. Voest, J. B. A. G. Haanen, T. N. Schumacher, Low and variable tumor reactivity of the intratumoral TCR repertoire in human cancers. Nature Medicine 25, 89–94 (2019).

42. N. Collins, X. Jiang, A. Zaid, B. L. Macleod, J. Li, C. O. Park, A. Haque, S. Bedoui, W. R. Heath, S. N. Mueller, T. S. Kupper, T. Gebhardt, F. R. Carbone, Skin CD4+ memory T cells exhibit combined cluster-mediated retention and equilibration with the circulation. Nat. Commun. 7, 11514 (2016).

43. T. N. Burn, J. Schröder, L. C. Gandolfo, M. Osman, E. N. Wainwright, E. Y. N. Lam, K. M. McDonald, R. B. Evans, S. Li, D. Rawlinson, L. Dryburgh, A. Zaid, Z. Maliga, D. Schienstock, P. Meiser, H. J. Lee, H. Lai, M. L. Moreira, P. Zareie, L. H.-Y. Lee, L. Huq, S. N. Christo, J. J. W. Seow, K. A. Ching, S. M. Guillaume, K. Knezevic, S. L. Park, M. Evrard, J. Waithman, T. Gebhardt, S. N. Mueller, G. E. Riddiough, M. V. Perini, S. C. H. Tsao, T. P. Speed, P. K. Sorger, S. Loi, F. R. Carbone, S. Gras, T. S. Fisher, B. J. Baaten, M. A. Dawson, L. K. Mackay, Antigen reactivity defines tissue-resident memory and exhausted T cells in tumours. bioRxiv, 2025.07.23.666465 (2025).

44. B. V. Kumar, W. Ma, M. Miron, T. Granot, R. S. Guyer, D. J. Carpenter, T. Senda, X. Sun, S.-H. Ho, H. Lerner, A. L. Friedman, Y. Shen, D. L. Farber, Human Tissue-Resident Memory T Cells Are Defined by Core Transcriptional and Functional Signatures in Lymphoid and Mucosal Sites. Cell reports 20, 2921–2934 (2017).

45. E. L.-H. Leung, R.-Z. Li, X.-X. Fan, L. Y. Wang, Y. Wang, Z. Jiang, J. Huang, H.-D. Pan, Y. Fan, H. Xu, F. Wang, H. Rui, P. Wong, H. Sumatoh, M. Fehlings, A. Nardin, P. Gavine, L. Zhou, Y. Cao, L. Liu, Longitudinal high-dimensional analysis identifies immune features associating with response to anti-PD-1 immunotherapy. Nat. Commun. 14, 5115 (2023).

46. T. Sato, Y. Ogawa, M. Kinoshita, Y. Nagasaka, S. Shimada, A. Momosawa, T. Kawamura, Preferential CD101 expression on resident memory regulatory T cells and IFN-γ-producing CD8+ resident memory T cells in human epidermis. J. Dermatol. Sci., doi: 10.1016/j.jdermsci.2025.07.002 (2025).

47. L. R. Soares, L. Tsavaler, A. Rivas, E. G. Engleman, V7 (CD101) ligation inhibits TCR/CD3-induced IL-2 production by blocking Ca2+ flux and nuclear factor of activated T cell nuclear translocation. J Immunol Baltim Md 1950 161, 209–17 (1998).

48. W. H. Hudson, J. Gensheimer, M. Hashimoto, A. Wieland, R. M. Valanparambil, P. Li, J.-X. Lin, B. T. Konieczny, S. J. Im, G. J. Freeman, W. J. Leonard, H. T. Kissick, R. Ahmed, Proliferating Transitory T Cells with an Effector-like Transcriptional Signature Emerge from PD-1+ Stem-like CD8+ T Cells during Chronic Infection. Immunity 51, 1043–1058.e4 (2019).

49. T. Duhen, R. Duhen, R. Montler, J. Moses, T. Moudgil, N. F. de Miranda, C. P. Goodall, T. C. Blair, B. A. Fox, J. E. McDermott, S.-C. Chang, G. Grunkemeier, R. Leidner, R. B. Bell, A. D. Weinberg, Co-expression of CD39 and CD103 identifies tumor-reactive CD8 T cells in human solid tumors. Nature Communications 9, 2724 (2018).

50. T. Hirai, Y. Yang, Y. Zenke, H. Li, V. K. Chaudhri, J. S. D. L. C. Diaz, P. Y. Zhou, B. A.-T. Nguyen, L. Bartholin, C. J. Workman, D. W. Griggs, D. A. A. Vignali, H. Singh, D. Masopust, D. H. Kaplan, Competition for Active TGFβ Cytokine Allows for Selective Retention of Antigen-Specific Tissue- Resident Memory T Cells in the Epidermal Niche. Immunity 54, 84–98.e5 (2021).

51. T. B. Shabaneh, A. Molodtsov, S. M. Steinberg, P. Zhang, G. M. Torres, G. A. Mohamed, A. Boni, T. J. Curiel, C. V. Angeles, M. J. Turk, Oncogenic BRAFV600E governs regulatory T cell recruitment during melanoma tumorigenesis. Cancer Res 78, canres.0365.2018 (2018).

52. J. N. Cohen, V. Gouirand, C. E. Macon, M. M. Lowe, I. C. Boothby, J. M. Moreau, I. K. Gratz, A. Stoecklinger, C. T. Weaver, A. H. Sharpe, R. R. Ricardo-Gonzalez, M. D. Rosenblum, Regulatory T cells in skin mediate immune privilege of the hair follicle stem cell niche. Sci. Immunol. 9, eadh0152 (2024).

53. R. S. Rodriguez, M. L. Pauli, I. M. Neuhaus, S. S. Yu, S. T. Arron, H. W. Harris, S. H.-Y. Yang, B. A. Anthony, F. M. Sverdrup, E. Krow-Lucal, T. C. MacKenzie, D. S. Johnson, E. H. Meyer, A. Löhr, A. Hsu, J. Koo, W. Liao, R. Gupta, M. G. Debbaneh, D. Butler, M. Huynh, E. C. Levin, A. Leon, W. Y. Hoffman, M. H. McGrath, M. D. Alvarado, C. H. Ludwig, H.-A. Truong, M. M. Maurano, I. K. Gratz, A. K. Abbas, M. D. Rosenblum, Memory regulatory T cells reside in human skin. J. Clin. Investig. 124, 1027–1036 (2014).

54. N. Ali, B. Zirak, R. S. Rodriguez, M. L. Pauli, H.-A. Truong, K. Lai, R. Ahn, K. Corbin, M. M. Lowe, T. C. Scharschmidt, K. Taravati, M. R. Tan, R. R. Ricardo-Gonzalez, A. Nosbaum, M. Bertolini, W. Liao, F. O. Nestle, R. Paus, G. Cotsarelis, A. K. Abbas, M. D. Rosenblum, Regulatory T Cells in Skin Facilitate Epithelial Stem Cell Differentiation. Cell 169, 1119–1129.e11 (2017).

